# Spatiotemporal Dynamics of Vascular Wall Cells during Ductus Arteriosus Development and Closure

**DOI:** 10.64898/2026.11.23.768583

**Authors:** Christine Sanderson, Irusha Dahal, Arnoneel Sinha, Martina Zoccheddu, Ashok Sharma, Jeff Reese, Elaine L. Shelton, James D. St. Louis, Brian K. Stansfield, Junichi Saito

## Abstract

**Background:** The transition from fetal to postnatal circulation requires closure of the ductus arteriosus (DA), involving smooth muscle cell contraction followed by anatomical remodeling. However, the spatial organization and developmental transcriptional dynamics of vascular wall cells during this transition remain poorly understood.

**Methods:** We analyzed vascular morphology and spatial transcript profiles of the mouse DA and aorta at five stages from embryonic day 14.5 (E14.5) to postnatal day 2.5 (P2.5), enabling paired comparisons within tissue sections. Selected findings were compared with an independent mouse single-cell RNA-sequencing dataset. Histological and spatial transcript analyses were also performed on human DA tissue and the adjacent pulmonary artery.

**Results:** The DA exhibited transiently higher cell-cycle-associated transcription across multiple vascular wall populations at E16.5, preceding medial thickening at E18.5 and postnatal luminal closure. Prenatally, DA mural cells exhibited greater contractile-associated gene expression than aortic mural cells. This difference diminished after birth, accompanied by a marked shift toward synthetic-like transcriptional programs associated with extracellular matrix remodeling and proteolysis. Ductal endothelial cells exhibited postnatal divergence from aortic endothelial cells in matrix- and adhesion-associated gene expression. Adventitial fibroblasts displayed distinct extracellular matrix- and insulin-like growth factor-associated expression profiles. In a human postnatal DA specimen, synthetic-like mural cells were enriched in regions of disrupted elastic lamellae and prominent collagen staining and shared selected remodeling-associated markers with postnatal mouse DA mural cells.

**Conclusions:** DA development and closure involve temporally organized transcriptional changes across vascular wall populations, characterized by prenatal cell-cycle-associated activity and rapid postnatal reorganization of mural transcriptional programs. These findings provide a spatial framework for understanding vascular cell plasticity and multicellular remodeling during DA closure.

**Clinical Implications:** The ductus arteriosus (DA) is a fetal vascular shunt that normally closes after birth. Persistent patency of the DA (patent ductus arteriosus [PDA]) is common in preterm infants, particularly those born extremely preterm, and hemodynamically significant PDA is associated with increased morbidity. While some infants require treatment to promote DA closure, patients with duct-dependent congenital heart disease require maintenance of DA patency to support pulmonary or systemic circulation. Current pharmacological approaches primarily regulate DA vascular tone through prostaglandin signaling rather than directly targeting the cellular mechanisms underlying vascular remodeling and permanent closure. Our findings identify temporally distinct smooth muscle cell transcriptional programs, accompanied by endothelial and fibroblast transcriptional changes, providing a cellular framework that may inform future therapeutic strategies to regulate DA remodeling and patency.

## INTRODUCTION

The ductus arteriosus (DA) is an essential fetal blood vessel that maintains fetal hemodynamics by diverting blood away from the nonventilated lungs into the systemic circulation through the descending aorta. Unlike other major thoracic arteries, the DA is developmentally programmed to close after birth and ultimately become a fibrous ligament. DA closure occurs in 2 phases. The first phase involves functional constriction, driven primarily by increased arterial oxygen tension and withdrawal of circulating prostaglandin E2. Subsequent anatomical remodeling transforms the constricted vessel into a fibrous ligament over the following weeks.^1–5^ Failure of these processes can result in patent ductus arteriosus (PDA), a common cardiovascular condition in newborns, particularly those born preterm.^2,6^

Smooth muscle cells (SMCs) are central to both phases of DA closure, although each phase requires distinct cellular behaviors. Functional constriction depends on the ability of medial SMCs to generate sufficient contractile force to narrow the vessel lumen, whereas anatomical closure involves SMC migration, extracellular matrix remodeling, and intimal thickening.^7–9^ These observations suggest that successful DA closure requires SMCs to support both contractile and matrix-remodeling programs during the perinatal period. Arterial SMCs are highly plastic and can transition between a differentiated contractile state and a synthetic-like state characterized by extracellular matrix production and remodeling.^10,11^ Such phenotypic transitions have been extensively investigated in vascular diseases, including atherosclerosis, restenosis, and aneurysm.^12–14^ In contrast, the DA provides a physiological developmental model in which SMC transcriptional programs can be examined across a defined perinatal time course and compared directly with neighboring arteries that remain patent.

Much of our understanding of the DA transcriptome has been derived from dissected tissues or dissociated cells. Bulk transcriptomic and laser-capture microdissection studies have identified differences in gene expression between the DA and aorta, including comparisons across rodent and human datasets.^15–17^ Human ductal SMCs have also been isolated and characterized for their responses to oxygen levels in culture,^18^ while gene expression has been compared between closing and patent human ductal tissues.^19^ More recently, single-cell RNA sequencing of the mouse DA and aorta across late fetal and early postnatal stages has defined major vascular cell populations and revealed cell type-specific transcriptional changes associated with DA closure.^20,21^ Several molecular regulators of DA development and closure have also been identified, including *PTGER4*, encoding the prostaglandin E2 receptor EP4, which contributes to prenatal DA patency and anatomical remodeling, and *TFAP2B*, pathogenic variants of which cause Char syndrome and are associated with familial nonsyndromic PDA.^22–28^ These genes belong to an expanding repertoire of molecular regulators implicated in persistent DA patency.^29,30^ However, existing transcriptomic studies have not simultaneously characterized gene expression at single-cell resolution while preserving the spatial organization of vascular wall cells throughout DA development and closure. Consequently, how SMC transcriptional programs and other vascular wall cell populations are spatially organized and undergo developmental changes during DA closure remains poorly understood.

Here we applied spatial molecular imaging^31^ to the mouse DA and adjacent ascending and descending aorta across five developmental stages, from embryonic day 14.5 (E14.5) to postnatal day 2.5 (P2.5), and to human DA tissue and the adjacent pulmonary artery. By examining the DA and both aortic regions within the same tissue sections, we enabled paired comparisons between closing and non-closing vessels under matched experimental conditions. We hypothesized that DA closure is accompanied by temporally coordinated changes in spatially organized vascular wall transcriptional programs, particularly those associated with SMC contraction and extracellular matrix remodeling. To test this hypothesis, we mapped vascular wall cell populations and their transcriptional programs *in situ* across development and investigated whether similar mural cell programs were detectable in human ductal tissue.

## RESULTS

### DA medial thickening precedes postnatal luminal closure

To characterize structural changes in the ductus arteriosus (DA) during perinatal development, we examined cross-sections of the DA and ascending aorta at E14.5, E16.5, E18.5, P0.5, and P2.5 by immunofluorescence staining for CD31, ACTA2, and DAPI. Representative images demonstrated progressive thickening of the DA media during prenatal development, followed by luminal closure after birth (Figure 1A). Quantitative morphometric analyses confirmed these changes (Figure 1B). At E14.5 and E16.5, medial wall thickness, medial wall area, and lumen area did not differ significantly between the DA and aorta. By E18.5, the DA exhibited significantly greater medial wall thickness than the aorta, a difference that persisted through P2.5. Whereas the aortic lumen progressively enlarged during development, the DA lumen narrowed and underwent closure after birth. Together, these findings demonstrate that DA remodeling begins prenatally with medial thickening, suggesting a potential role for smooth muscle cells (SMCs) in the structural changes preceding postnatal luminal closure.

**Figure 1.**
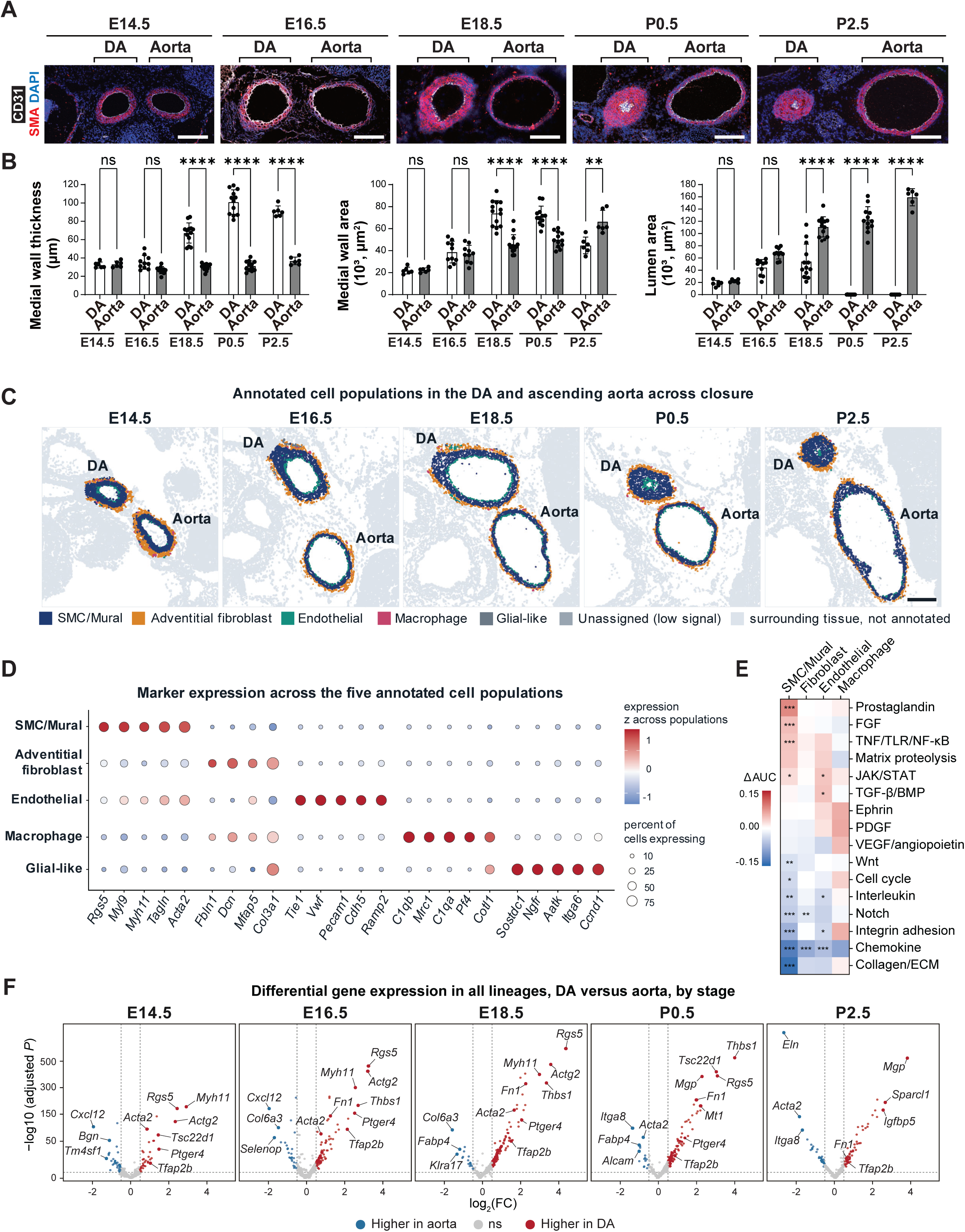
Structural changes, cellular composition, and transcriptional profiles of the mouse ductus arteriosus and adjacent aortas during perinatal development. **(A)** Representative immunofluorescence images of the ductus arteriosus (DA) and ascending aorta at E14.5, E16.5, E18.5, P0.5, and P2.5, showing CD31 (white), ACTA2 (red), and DAPI (blue). Scale bar, 100 μm. **(B)** Quantification of medial wall thickness, medial wall area, and lumen area in the DA and ascending aorta across developmental stages. n = 6, 10, 13, 12, and 6 animals at E14.5, E16.5, E18.5, P0.5, and P2.5, respectively. ns, not significant; \*\**P* < 0.01; \*\*\*\**P* < 0.0001, by one-way ANOVA followed by Tukey’s multiple-comparison test. **(C)** Spatial distribution of annotated cell populations in representative tissue sections from each developmental stage. Pale gray denotes surrounding unannotated tissue. Scale bar, 200 μm. **(D)** Expression of selected marker genes across the five annotated cell populations. Dot size represents the percentage of cells expressing each gene, and color indicates scaled expression levels. **(E)** Differences in signaling-module scores between the DA and pooled aorta across annotated cell populations. Red indicates higher module scores in the DA, whereas blue indicates higher scores in the aorta. The color scale is capped at ±0.15 ΔAUC. n = 20 sections; macrophages were excluded from statistical testing. For each module and cell population, within-section DA–aorta differences were tested against zero using a one-sample *t* test, followed by Benjamini-Hochberg correction across the 16 modules within each population. \**q* < 0.05; \*\**q* < 0.01; \*\*\**q* < 0.001. **(F)** Stage-specific differential gene expression between the DA and pooled aorta. The y axis represents −log_10_ of the adjusted *P* value. Red indicates higher expression in the DA, blue indicates higher expression in the aorta, and gray indicates genes without statistically significant differences. Selected genes are highlighted and labeled. These cell-level results were used to screen candidate genes; inferential analyses used section-level comparisons. AUC, area under the curve; BMP, bone morphogenetic protein; DA, ductus arteriosus; ECM, extracellular matrix; FGF, fibroblast growth factor; JAK, Janus kinase; NF-κB, nuclear factor κB; PDGF, platelet-derived growth factor; SMC, smooth muscle cell; STAT, signal transducer and activator of transcription; TGF-β, transforming growth factor β; TLR, Toll-like receptor; TNF, tumor necrosis factor; VEGF, vascular endothelial growth factor.

### Spatial atlas of the DA and aorta reveals cell population-specific transcriptional identities

To characterize the cellular composition and spatial organization of the vessel wall, we annotated 48,822 vessel-wall cells across 20 tissue sections from four animals per developmental stage and classified them into five major populations: mural cells, adventitial fibroblasts, endothelial cells, macrophages, and glial-like cells (Figure 1C). These populations exhibited distinct spatial distributions, with mural cells predominantly localized to the tunica media, adventitial fibroblasts to the outer vessel wall, and endothelial cells lining the lumen (Supplemental Figure S1A). Cell identities were further supported by canonical marker gene expression. One-versus-rest differential expression analysis using the 965-gene panel identified *Acta2* and *Tagln* as markers of mural cells; *Col3a1* and *Col1a1* of adventitial fibroblasts; *Vwf* and *Pecam1* of endothelial cells; *C1qb* and *C1qa* of macrophages; and *Itga6* and *Ngfr* for glial-like cells (Figure 1D). The glial-like population was localized to the outer boundary of the adventitia, consistent with previous single-cell studies identifying Schwann cells in the aorta and nerve-associated cells in the DA and other great vessels.^20,32,33^ However, the commercial targeted gene panel lacked sufficient lineage-specific markers to characterize this population more precisely, and glial-like cells were therefore excluded from subsequent transcriptional analyses. Although *Notch3* and *Pdgfrb* were included in the panel, their expression did not reliably distinguish pericytes from other mural cells. Accordingly, SMCs and other mural cells were analyzed as a single population. The integrated UMAP embedding revealed distinct transcriptional identities among the major vascular wall cell populations and differences associated with vessel of origin, whereas cells from the ascending and descending aorta and across developmental stages showed substantial overlap (Supplemental Figure S1B).

Mural cells constituted the predominant population in both vessels across all developmental stages, based on the relative proportions of the five annotated cell populations (Supplemental Figure S1C). During prenatal development, the DA exhibited a higher proportion of mural cells than the paired aorta (67.2% versus 62.2% at E14.5, 66.7% versus 59.5% at E16.5, and 75.0% versus 65.5% at E18.5; *P* = 0.021, 0.002, and 0.0001, respectively), accompanied by a lower proportion of adventitial fibroblasts (22.1% versus 26.1%, 20.3% versus 24.8%, and 16.3% versus 22.0%, respectively; *P* = 0.020, 0.014, and 0.004). The higher mural cell proportion in the DA persisted at P0.5 (75.2% versus 71.2%; *P* = 0.046), but the difference was no longer detected at P2.5. At P2.5, the fibroblast proportion was higher in the DA than in the aorta (21.1% versus 16.1%; *P* = 0.009). The endothelial cell proportion was lower in the DA at E16.5 (10.1% versus 14.4%; *P* = 0.004) and P0.5 (6.1% versus 10.6%; *P* = 0.004). The macrophage proportion was higher in the DA at E14.5 (3.0% versus 1.3%; *P* = 0.0004) but lower at E18.5 (0.7% versus 2.2%; *P* = 0.001).

Together, these findings establish a spatially resolved atlas of the developing DA and aorta, revealing distinct cell population-specific transcriptional identities and developmental differences in the relative composition of vessel-wall cells.

### Transcriptional differences between the DA and aorta

The ascending and descending aortas were pooled as a single comparator because only 17 of 963 genes were differentially expressed between these two regions, compared with 218 genes differentially expressed between the DA and pooled aorta. Consistent with this transcriptional similarity, cells from the two aortic segments showed substantial overlap in the integrated UMAP embedding (Supplemental Figure S1B). The 17 segment-specific genes were predominantly extracellular matrix-associated genes, including *Col3a1*, *Eln*, and *Bgn*, in line with the regional differences in embryonic aortic gene expression reported previously.^34^ When the DA was compared with each aortic segment separately, 177 and 201 genes differed from the ascending and descending aorta, respectively, and 126 genes were shared with concordant direction.

To further characterize vessel-specific transcriptional differences, we calculated scores for 16 gene panel-restricted signaling modules and compared the DA with the pooled aorta within each section for each population (Figure 1E). Most differences in signaling-module scores were observed in mural cells, with 11 of the 16 modules differing between vessels. The prostaglandin signaling module exhibited the strongest enrichment in the DA (*q* = 0.0003), with a significant difference detected only in mural cells. The FGF and TNF/TLR/NF-κB signaling modules also differed between vessels exclusively in mural cells. Fewer differences were observed in endothelial cells and adventitial fibroblasts. The JAK/STAT signaling module showed higher scores in the DA in both mural cells (*q* = 0.023) and endothelial cells (*q* = 0.038), whereas the TGF-β/BMP signaling module showed higher scores in DA endothelial cells (*q* = 0.038).

To identify vessel-specific transcriptional differences across development, we compared gene expression between the DA and pooled aorta at each developmental stage (Figure 1F). The number of differentially expressed genes increased from 58 at E14.5 to a maximum of 164 at E18.5, before declining to 97 at P2.5, suggesting that the greatest number of detected vessel-specific transcriptional differences occurred during late gestation. The composition of these transcriptional signatures also changed across development. During prenatal development, DA-enriched genes included contractile-associated genes such as *Myh11*, *Actg2*, and *Rgs5*, whereas postnatal DA-enriched genes included matricellular and stress-associated genes such as *Mgp*, *Thbs1*, *Tsc22d1*, and *Mt1*. Nine genes (*Cald1*, *Col6a2*, *Hspb1*, *Mmp14*, *Sox9*, *Tfap2b*, *Thbs1*, *Tsc22d1*, and *Tshz2*) were consistently enriched in the DA across all five developmental stages, whereas only *Cxcl12* was consistently enriched in the aorta. In the independent scRNA-seq dataset GSE188202, eight of these nine genes were enriched in DA SMCs relative to aortic SMCs at E18.5, P0.5, and P5, and *Cxcl12* was enriched in aortic SMCs at all three ages (Supplemental Table S1). These findings indicate that the DA maintains a core transcriptional signature while exhibiting developmental changes in its vessel-specific gene-expression profile. Notably, two established DA-associated genes exhibited distinct temporal patterns: *Ptger4* remained enriched in the DA through P0.5 but not at P2.5, whereas *Tfap2b* remained DA-enriched throughout all five stages.

### Developmental transitions in mural cell transcriptional programs

To investigate the spatial and transcriptional heterogeneity of the mural compartment, we classified mural cells into five transcriptional programs using curated marker sets from the targeted gene panel (Figures 2A and 2B; Supplemental Figure S1D): contractile (e.g., *Myh11* and *Acta2*), proliferative (e.g., *Mki67* and *Pcna*), contractile-remodeling (e.g., *Igf2* and *Thbs1*), matricellular-remodeling (e.g., *Fn1* and *Bgn*), and matrix-remodeling (e.g., *Mmp2* and *Mmp9*).

**Figure 2.**
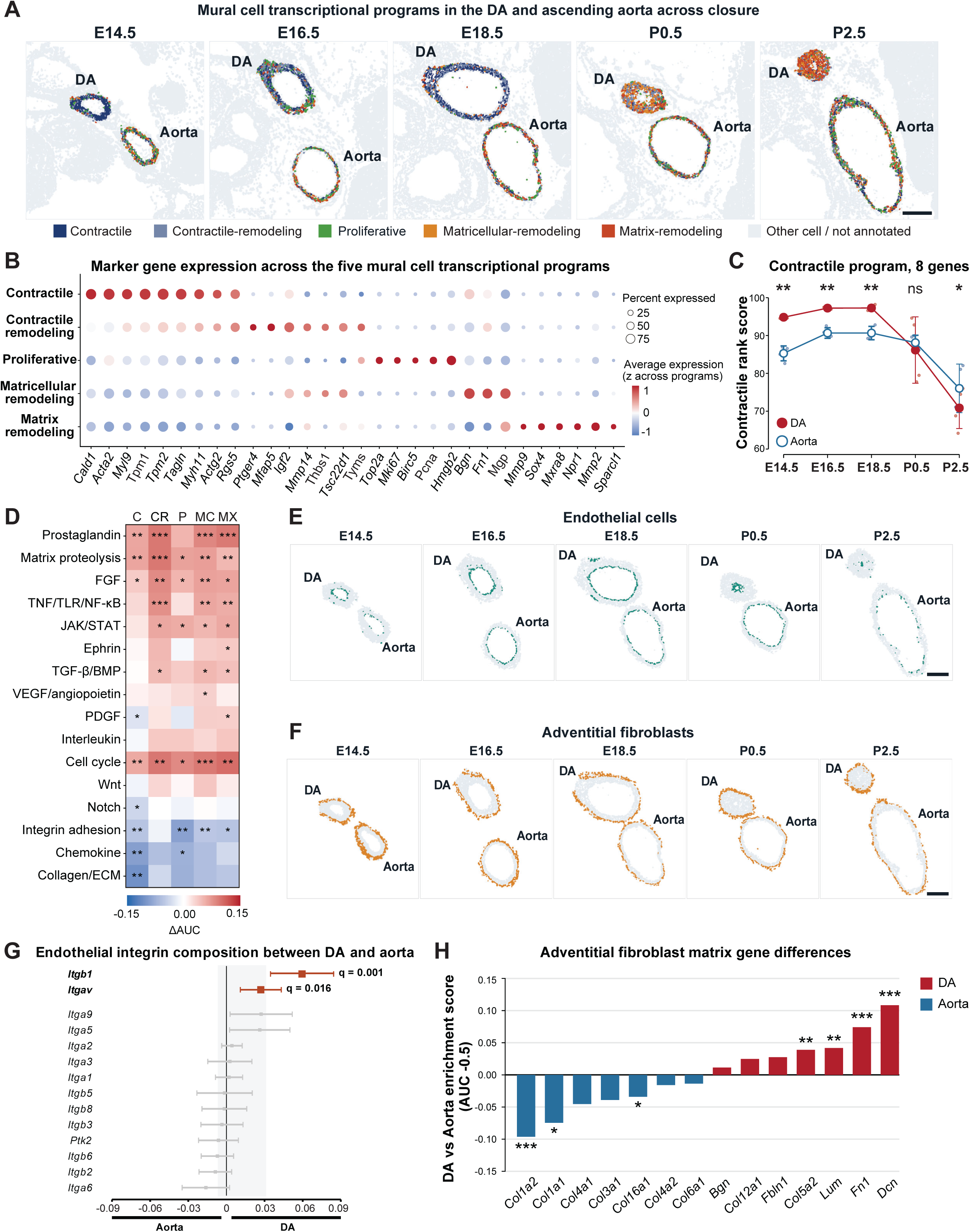
Mural cell transcriptional programs, endothelial integrins, and adventitial fibroblast matrix genes in the ductus arteriosus and aorta. **(A)** Spatial distribution of mural cell transcriptional programs in the ductus arteriosus (DA) and ascending aorta across developmental stages. Nonmural cells are shown in gray. Scale bar, 200 μm. **(B)** Expression of selected marker genes across the five mural cell transcriptional programs. Dot size represents the percentage of cells expressing each gene, and color indicates scaled expression levels. **(C)** Contractile rank scores of mural cells in the DA and pooled aorta across developmental stages. Scores represent the mean percentile rank of eight contractile-associated genes among all panel genes within each cell (range, 0–100). Small points represent individual tissue sections, whereas larger symbols and error bars indicate the mean ± SD across sections (n = 4 sections per stage). Paired two-sided *t* tests were performed at each stage, followed by Benjamini-Hochberg correction across the five stages. ns, not significant; \**q* < 0.05; \*\**q* < 0.01. **(D)** Differences in signaling-module scores between DA and aortic mural cells within each transcriptional program. Red indicates higher module scores in the DA, whereas blue indicates higher scores in the aorta. Programs are abbreviated as follows: C, contractile; CR, contractile-remodeling; P, proliferative; MC, matricellular-remodeling; and MX, matrix-remodeling. Signaling-module abbreviations are defined in Figure 1. \**q* < 0.05; \*\**q* < 0.01; \*\*\**q* < 0.001. **(E)** Spatial distribution of endothelial cells across developmental stages. Scale bar, 200 μm. (F) Spatial distribution of adventitial fibroblasts across developmental stages. Scale bar, 200 μm. **(G)** Differences in expression of integrin adhesion genes between DA and aortic endothelial cells (14 genes). Points represent the mean DA–aorta expression contrast across 20 paired sections, and error bars indicate 95% confidence intervals. Red indicates higher expression in the DA, blue indicates higher expression in the aorta, and gray indicates genes without statistically significant differences. The shaded region represents the 10th–90th percentile range of the corresponding contrasts for 300 randomly selected panel genes. Adjusted *q* values are displayed for genes with statistically significant differences. (H) Differences in expression of selected extracellular matrix genes between DA and aortic adventitial fibroblasts. Red indicates higher expression in the DA, whereas blue indicates higher expression in the aorta (n = 20 sections). \**q* < 0.05; \*\**q* < 0.01; \*\*\**q* < 0.001. AUC, area under the curve; DA, ductus arteriosus; SD, standard deviation.

These programs were intended to characterize transcriptional heterogeneity rather than define discrete mural cell subtypes. Consistent with this approach, mural cells assigned to different programs showed substantial overlap in the integrated UMAP embedding rather than forming clearly separated clusters (Supplemental Figure S1E). Additional analyses provided limited evidence for discrete transcriptional states. Within each developmental stage, Hartigan’s dip tests were nonsignificant (*P* = 0.449–0.996), a two-component model was not favored at four of the five stages, and no individual marker distinguished a program with an area under the receiver operating characteristic curve (AUC) greater than 0.68. Together, these findings suggest that mural transcriptional programs represent partially overlapping patterns of contractile- and remodeling-associated gene expression rather than fixed cellular identities, consistent with SMC phenotypic plasticity.

To quantify contractile-associated gene expression independently of program assignment, we calculated a rank-based composite score using eight contractile-associated genes (Supplemental Table S2) within each mural cell (Figure 2C). Contractile scores were significantly higher in DA mural cells than in aortic mural cells at E14.5 (93.4 versus 84.6; *q* = 0.0018), E16.5 (97.1 versus 90.6; *q* = 0.0018), and E18.5 (96.8 versus 89.9; *q* = 0.0018). This difference was no longer detected at P0.5 (86.2 versus 87.9; *q* = 0.67). By P2.5, contractile scores were significantly lower in the DA than in the aorta (70.8 versus 76.0; *q* = 0.019). These findings demonstrate that prenatal enrichment of contractile-associated gene expression in DA mural cells diminishes after birth and reverses by P2.5.

The distribution of mural cell transcriptional programs also changed markedly during DA development (Supplemental Figure S1F). In the DA, the proportion of mural cells assigned to the contractile program decreased from 61.2% at E14.5 to 4.7% at P2.5, whereas the proportion assigned to the matrix-remodeling program increased from 7.4% to 55.4%. The matricellular-remodeling program reached its highest proportion at P0.5 (37.8%) and remained prominent at P2.5 (26.0%). In contrast, the aortic mural cell compartment exhibited relatively stable program composition, with contractile-program cells accounting for 27.4% at both E14.5 and P2.5 and matrix-remodeling-program cells accounting for 17.6% and 19.9%, respectively. Postnatally, particularly at P2.5, the DA contained a significantly lower proportion of contractile-program cells and a higher proportion of matrix-remodeling-program cells than the aorta (*P* < 0.0001 for both). Consistent with these findings, reanalysis of an independent published mouse single-cell RNA-sequencing dataset revealed lower contractile-associated gene expression and higher expression of *Klf4* and matrix-remodeling-associated genes in postnatal DA SMCs relative to aortic SMCs (Supplemental Table S1).^20^

### Vessel-specific signaling differences persist within matched mural cell programs

The DA and aorta differed in the relative proportions of the five mural cell programs, so the vessel-specific differences in signaling-module scores (Figure 1E) could simply reflect that difference in composition rather than differences within corresponding programs. To test this, we compared signaling-module scores between the DA and aorta separately within each mural cell program (Figure 2D). Of the 80 program-by-module comparisons, 34 met the significance criteria under both scoring specifications, with concordant directions of difference. The cell cycle, FGF, matrix-proteolysis, and prostaglandin signaling modules showed higher scores in the DA in four of the five mural programs, whereas the JAK/STAT and TNF/TLR/NF-κB modules showed higher scores in three programs and the TGF-β/BMP module in two. Conversely, the integrin-adhesion module showed higher scores in the aorta in four programs. No module exhibited significant differences in opposite directions across programs. Notably, 29 of the 34 significant comparisons were directionally concordant with the corresponding whole-compartment analyses. At the gene level, *Itga8* was the most aorta-enriched integrin in mural cells, with higher expression in aortic than in ductal mural cells in all 20 sections (section-level mean AUC − 0.5 = −0.11, *P* < 0.001). In GSE188202, *Itga8* expression in aortic SMCs was two- to four-fold the ductal level at E18.5, P0.5, and P5 (Supplemental Table S1). Together, these findings indicate that vessel-specific differences in signaling-module scores persist within corresponding mural cell programs and cannot be explained solely by differences in their relative proportions.

### Transient prenatal cell-cycle-associated transcriptional activity

The DA undergoes medial thickening during prenatal development (Figure 1A). To investigate whether this process was associated with developmental changes in cell-cycle-associated transcriptional activity, we analyzed cell-cycle module scores across the major structural cell populations of the vascular wall. When scores were aggregated across endothelial cells, mural cells, and adventitial fibroblasts, the DA exhibited significantly higher cell-cycle module scores than the paired aorta only at E16.5 (*q* = 0.035), with no significant differences detected at the other developmental stages (Supplemental Figure S2A and S2B). At E16.5, endothelial cell-cycle module scores were approximately 41% higher in the DA than in the aorta. Higher scores were also observed in DA mural cells and adventitial fibroblasts, with increases of approximately 13% and 29%, respectively. Thus, the transient E16.5 increase in cell-cycle-associated transcription extended across the major structural cell populations of the DA wall, with the largest relative difference observed in endothelial cells.

### Endothelial adhesion and remodeling genes differ between the vessels

Endothelial cells lined the lumen, whereas adventitial fibroblasts occupied the outer vessel wall in both vessels throughout development (Figures 2E and 2F). To compare adhesion-related gene expression between ductal and aortic endothelial cells, we examined the genes comprising the integrin-adhesion module (Figure 2G). *Itgb1* and *Itgav* were enriched in DA endothelial cells. Despite these gene-specific differences, the 14 integrin-adhesion genes as a set did not exhibit a significant shift relative to a background of 300 genes from the targeted panel (*P* = 0.38), indicating selective differences in integrin-subunit expression rather than coordinated enrichment across the gene set.

To identify endothelial genes exhibiting stage-dependent differences between the DA and aorta, we ranked all detected endothelial genes according to the strength of their vessel-by-stage interactions. *Mgp* ranked first (*q* = 0.0001), followed by *Igfbp7* and *Vim* (*q* = 0.0003 each). The vessel-specific difference in *Vim* expression was greatest postnatally and remained elevated at P2.5 (Supplemental Figure S2C). Other highly ranked genes included *Lmna*, *Hsp90ab1*, *Hspb1*, *Col1a2*, *Col5a2*, *Col1a1*, *Col3a1*, and *Clu*, indicating that stage-dependent vessel differences extended beyond canonical endothelial markers to include several extracellular matrix-associated genes. Among the 27 endothelial genes with significant stage-dependent vessel differences (*q* < 0.05), 12 showed little difference between vessels through E18.5 and diverged after birth (Supplemental Methods).

Comparison with the independent scRNA-seq dataset GSE188202 provided additional gene-level evidence of vessel-specific endothelial expression differences after birth.^20^ The strongest temporal agreement was observed for *Vim*, which became enriched in DA endothelial cells at P0.5 in the external dataset. In contrast, *Col5a2* exhibited postnatal enrichment in aortic endothelial cells*. Clu* and *Mgp* also showed persistent vessel-specific expression differences detectable before birth (Supplemental Table S1). Together, these findings provide independent directional support for the endothelial expression patterns identified by spatial transcript profiling.

### Ductal and aortic fibroblasts differ in matrix-associated gene expression and IGF-axis composition

To compare extracellular matrix-associated gene expression between ductal and aortic adventitial fibroblasts, we analyzed genes represented in the targeted panel across all 20 tissue sections. DA adventitial fibroblasts exhibited higher expression of *Dcn*, *Fn1*, *Lum*, and *Col5a2* than their paired aortic counterparts, whereas aortic fibroblasts showed higher expression of *Col1a1*, *Col1a2*, and *Col16a1* (*q* < 0.05; Figure 2H). We next investigated whether vessel-specific differences in adventitial fibroblast gene expression varied across developmental stages. Three genes (*Eln*, *Mmp2*, and *Cxcl12*) met the significance criterion (*q* < 0.05) and exhibited pronounced developmental changes in their vessel-specific expression differences (Supplemental Figure S2D). *Eln* expression was higher in DA fibroblasts than in paired aortic fibroblasts at E18.5, but this relationship reversed by P2.5. In contrast, *Mmp2* showed little vessel-specific difference prenatally but became enriched in DA fibroblasts after birth.

To investigate whether growth factor-related gene expression also differed between vessels, we examined genes associated with the insulin-like growth factor (IGF) axis in adventitial fibroblasts (Supplemental Figure S2E). Across developmental stages, DA fibroblasts exhibited higher expression of *Igf1* and *Igf2*, whereas aortic fibroblasts showed higher expression of *Igfbp5* and *Igf2r*, with *Igf2* exhibiting the largest vessel-specific difference. However, the IGF-axis gene set as a whole did not show a significant shift between vessels, indicating gene-specific differences rather than coordinated enrichment across the gene set. The vessel-specific difference in *Igfbp5* was most pronounced at E16.5 (Supplemental Figure S2F), coinciding with increased prenatal cell-cycle-associated transcription, whereas the *Igf2* difference was greatest at P2.5. Together, these findings reveal distinct patterns of IGF-axis gene expression between ductal and aortic adventitial fibroblasts, without evidence of coordinated enrichment across the IGF-axis gene set.

### Histological and spatial gene expression analysis of human DA tissue

Finally, we examined human DA tissue and the adjacent pulmonary artery to investigate whether spatial and transcriptional features observed in mice were also detectable in human tissue. Elastica van Gieson and Masson’s trichrome staining of paraffin-embedded sections revealed disruption of the elastic lamellar architecture in the ductal region, accompanied by more prominent collagen staining (Supplemental Figure S3A–D). In contrast, the pulmonary artery exhibited relatively preserved elastic lamellae and less prominent collagen staining.

Of the 69,390 quality-controlled cells, 24,858 unassigned or VWF-high low-count cells and 3,767 erythrocytes and hemoglobin-associated low-count cells were set aside, and the remaining 40,765 cells were classified into 11 annotated cell populations and transcriptional programs based on their expression profiles. Within the mural cell compartment, three prominent transcriptional programs were identified: synthetic-like (18.8%), contractile (15.1%), and stress-response (12.2%), with percentages calculated relative to all annotated cells (Supplemental Figure S3E and S3F). The stress-response program was characterized by immediate-early genes (*FOS, JUN,* and *JUNB*) and may reflect tissue collection or processing conditions rather than a distinct developmental state. These programs exhibited partially distinct spatial distributions within the specimen (Supplemental Figure S3G). Notably, cells assigned to the synthetic-like program were enriched in the ductal region, spatially coinciding with areas of disrupted elastic lamellae and more prominent collagen staining.

Despite their spatial differences, the three mural transcriptional programs occupied adjacent and partially overlapping regions of the integrated transcriptional embedding rather than forming completely separated clusters (Supplemental Figure S3H). Nevertheless, each program exhibited a distinguishable marker-expression profile (Supplemental Figure S3I), supporting their interpretation as partially overlapping transcriptional programs rather than discrete mural cell identities.

To explore potential cross-species similarities, we compared the gene-expression signatures of postnatal mouse DA mural cells and human synthetic-like mural cells using genes shared between the targeted panels. This analysis identified 12 shared markers, including *COL6A2*, *MMP2*, *VCAN*, *TSC22D1*, and *MGP* (*q* < 0.05; Supplemental Figure S3J). Although descriptive, these findings suggest shared remodeling-associated transcriptional features between postnatal mouse and human DA mural cells.

## DISCUSSION

The DA undergoes a remarkable developmental transition, maintaining patency throughout fetal life before rapidly remodeling to achieve permanent closure after birth. Although the physiological mechanisms regulating ductal constriction have been extensively investigated, how the DA establishes its specialized vascular wall during prenatal development and subsequently undergoes postnatal remodeling remains poorly understood.^1–3,6–9^ In this study, we used spatial transcript profiling across five developmental stages to characterize the cellular and transcriptional dynamics of the mouse DA relative to the aorta. Our findings reveal a temporally organized sequence of vascular wall changes, including a transient prenatal increase in cell-cycle-associated transcriptional activity, developmental changes of contractile SMC programs, and a pronounced postnatal shift toward remodeling-associated mural cell states. Together, these findings suggest that prenatal DA development and postnatal remodeling involve distinct but temporally coordinated transcriptional programs, providing a framework for understanding the relationship between perinatal vascular wall maturation and postnatal DA closure.

One of our notable findings was the transient increase in cell-cycle-associated transcriptional activity in the DA at E16.5, preceding prenatal medial wall thickening. Medial thickening is the earliest structural difference between the DA and aorta in this series (Figure 1B), and whether it arises from cell proliferation, cell enlargement, or recruitment of cells into the wall has not been resolved. A wall-wide increase in cell-cycle-associated transcription two days before we first detected medial thickening is consistent with a proliferative contribution during this window, although cell-cycle transcript abundance indicates entry into the cell cycle rather than completed cell division. This increase was observed across mural cells, endothelial cells, and adventitial fibroblasts, suggesting that prenatal DA wall development involves multiple cellular compartments rather than mural cells alone. Cell-cycle-associated transcriptional activity was also detected within contractile mural cells, suggesting that prenatal wall growth may occur within a predominantly contractile mural compartment before its transcriptional composition changes following birth. Although previous scRNA-seq analysis identified increased cell-cycle-associated expression in postnatal DA cells at P0.5, which diminished by P5,^20^ our spatial analysis detected significant DA-enriched wall-wide cell-cycle activity only at E16.5. These observations may reflect differences in developmental timing, sampling resolution, or analytical approaches and raise the possibility that cell proliferation occurs during more than one developmental interval.

The developmental redistribution of mural transcriptional programs represents another major finding of this study. During prenatal development, DA mural cells were predominantly associated with contractile programs, whereas postnatal mural cells exhibited increased representation of matricellular- and matrix-remodeling programs. This transition resembles the contractile-to-synthetic phenotypic modulation described in vascular diseases,^10,11^ but occurs in the DA as a part of a physiological developmental process. Importantly, the five mural programs were not completely segregated in transcriptional or anatomical space, suggesting that they represent overlapping transcriptional states rather than discrete cell identities. A previous scRNA-seq study similarly reported reduced contractile gene expression and increased remodeling-associated transcripts in postnatal ductal SMCs, although individual gene-level trajectories were not uniformly concordant.^20^ Our spatial analysis extends these observations by preserving the anatomical context of mural cell states and demonstrating their developmental redistribution within the vessel wall.

The postnatal enrichment of remodeling-associated mural programs is consistent with the extensive structural reorganization required for anatomical DA closure. Previous histological studies have demonstrated that anatomical closure involves changes in elastic architecture, mural cell migration, and extracellular matrix remodeling.^7,9,35^ In our study, matrix-remodeling and proteolysis-associated transcripts were enriched in postnatal ductal mural cells, supporting a relationship between the observed transcriptional changes and these established structural processes. Our analysis of a human DA specimen provided additional descriptive evidence of remodeling-associated mural states. Synthetic-like SMCs were enriched in a ductal region characterized by disrupted elastic lamellae and increased collagen deposition, and several of their markers overlapped with those enriched in postnatal mouse DA mural cells. These observations are consistent with previous human and cross-species studies identifying SMC- and matrix-associated differences in DA tissues.^17–19^ Although the human findings are limited to a single specimen obtained after clinical interventions, they provide a preliminary basis for investigating whether similar remodeling-associated mural programs are present across human DA specimens.

Beyond mural cells, adventitial fibroblasts exhibited distinct transcriptional profiles between the DA and aorta, particularly in genes associated with extracellular matrix organization. Ductal fibroblasts showed higher expression of *Dcn*, *Fn1*, *Lum*, and *Col5a2*, whereas aortic fibroblasts exhibited higher expression of *Col1a2*, *Col1a1*, and *Col16a1*. These findings suggest vessel-specific differences in extracellular matrix composition rather than a generalized increase in matrix production. Notably, *Mmp2* became enriched in ductal fibroblasts at P2.5, consistent with previous evidence implicating MMP2 in structural remodeling of the DA.^35^ We also identified vessel-specific differences in the IGF axis, with higher *Igf1* and *Igf2* expression in ductal fibroblasts and higher *Igfbp5* and *Igf2r* expression in aortic fibroblasts. Although these differences did not represent coordinated enrichment of the IGF axis gene set, they suggest distinct growth factor–associated transcriptional environments between the vessels. Together, these findings highlight the potential contribution of adventitial fibroblasts to the extracellular matrix environment accompanying DA development and postnatal remodeling.

Endothelial cells also exhibited distinct developmental transcriptional changes between the DA and aorta. Among genes showing significant stage-dependent vessel differences, *Vim* displayed particularly pronounced divergence between the two vessels during the perinatal period. This finding is consistent with a previous study that identified developmental changes in endothelial *Vim* expression and demonstrated incomplete postnatal DA closure in global *Vim*-knockout mice.^20^ Although these findings suggest a potential role for vimentin in DA remodeling, its endothelial-specific contribution remains to be established. We also observed differential expression of integrin subunits, including *Itgb1* and *Itgav*, suggesting differences in endothelial adhesion-receptor composition rather than generalized activation of integrin signaling. These differences may influence endothelial interactions with the subendothelial extracellular matrix, consistent with previous observations of integrin-associated changes during primate DA closure.^36^ Together, these findings suggest that endothelial transcriptional remodeling accompanies the perinatal transition of the DA and may contribute to changes in the vascular wall environment during anatomical closure.

Several limitations should be considered when interpreting our findings. First, mural cells were analyzed as a single compartment because *Notch3* and *Pdgfrb* did not reliably distinguish pericytes from SMCs using the targeted panel. Moreover, the five mural transcriptional programs represent overlapping patterns of gene expression rather than discrete cell types, and their relative proportions depend on the curated marker sets used for classification. Second, the 965-gene panel excludes potentially important regulators and limits comprehensive pathway analysis. Module scores reflect coordinated transcript abundance but do not directly measure protein activity, ligand–receptor engagements, or signaling flux. Third, because different animals were examined at each developmental stage, individual cells could not be tracked over time. Thus, the observed redistribution of mural transcriptional programs may reflect differential proliferation, survival, or cell recruitment in addition to transcriptional switching. Fourth, most imaging runs contained only one developmental stage, limiting the ability to distinguish developmental effects from imaging-run effects. Accordingly, our interpretations rely primarily on within-section comparisons between the DA and aorta, which were imaged together. Hemodynamic and structural differences between the vessels may also contribute to their transcriptional divergence. Finally, the external scRNA-seq dataset has limited biological replication for formal stage-specific validation, and the human analysis was restricted to a single specimen, reflecting the scarcity of such tissue, and the artery available for comparison in that specimen was the pulmonary artery rather than the aorta. In addition, transcriptomic cell-cycle signatures do not directly establish proliferation, and differences between datasets preclude definitive conclusions regarding the timing of cell expansion. Despite these limitations, the paired within-section design and the directional agreement of several findings with an independent single-cell dataset provide support for the principal observations.

Together, our findings reveal temporally organized changes in vascular wall cell states during DA development and closure. Prenatal increases in cell-cycle-associated transcriptional activity are followed by a postnatal shift from contractile-dominant to remodeling-associated mural cell states, accompanied by transcriptional changes in endothelial cells and adventitial fibroblasts. By resolving these changes within the intact vessel wall, our study highlights the multicellular nature of DA remodeling and identifies the DA as a valuable physiological model for investigating vascular SMC plasticity. Future studies incorporating whole-transcriptome spatial profiling, denser developmental sampling, pulse-labeling and lineage tracing, functional perturbation, and additional human specimens will be needed to determine the timing of proliferation in specific vascular wall populations, clarify the mechanisms underlying these developmental changes, and define their contributions to DA closure. Understanding the temporal regulation of these programs may provide new insights into DA development and vascular diseases characterized by altered SMC states.

## MATERIALS AND METHODS

### Study design and tissue collection

Two independent cohorts of C57BL/6J mice were used for immunofluorescence-based wall morphometry (47 animals) and CosMx spatial molecular imaging (20 animals). For morphometric analyses, one tissue section per animal was examined at five developmental stages: embryonic day (E) 14.5, E16.5, and E18.5 and postnatal day (P) 0.5 and P2.5, with at least six animals per developmental stage. For CosMx SMI profiling, 20 tissue sections from 20 distinct embryos or pups were analyzed, with four animals from more than one litter examined at each developmental stage. E0.5 was defined as noon on the day a vaginal plug was detected. Sex was not determined in mouse embryos or pups. Each section used for CosMx SMI profiling contained the ductus arteriosus (DA), ascending aorta, and descending aorta, enabling paired within-section comparisons among vessels. Cells from the ascending and descending aorta were pooled for transcriptomic comparisons between the DA and aorta, because the two segments differed in only 17 of 963 genes and overlapped in the integrated embedding (Results).

Human DA and adjacent pulmonary arterial tissue were obtained from a single patient with hypoplastic left heart syndrome at six months of age during cardiac surgery at Children’s Hospital of Georgia. The patient was born at 39 weeks of gestation with a birth weight of 4,180 g and received prostaglandin E1 during the first week after birth. The DA was subsequently stented and remained patent until surgery. The stent was removed before sample processing.

All mouse experiments were approved by the Institutional Animal Care and Use Committee at Augusta University (protocol 2025-1152) and performed in accordance with the NIH Guide for the Care and Use of Laboratory Animals. The use of de-identified human tissue was reviewed by the Institutional Review Board at Augusta University and determined not to constitute human subjects research.

### Immunofluorescence and wall morphometry

Mouse embryos and pups were fixed in 4% paraformaldehyde for 2 hours at 4°C, cryoprotected in 30% sucrose overnight, and embedded in Optimal Cutting Temperature (OCT) compound (Sakura Tissue-Tek). Transverse cryosections were cut at 5 μm from the junction of the descending aorta with the DA to the pulmonary artery and stored at −80°C until processing.

Sections were blocked and permeabilized in PBS containing 5% goat serum and 0.5% Triton X-100, followed by overnight incubation at 4°C with rat anti-mouse CD31 antibody (clone MEC 13.3; BD Pharmingen, 553370; 1:250). The following day, sections were incubated for 1 hour at room temperature in the dark with Alexa Fluor 488-conjugated goat anti-rat IgG (Invitrogen, A-11006; 1:500), Cy3-conjugated mouse monoclonal anti-α-smooth muscle actin antibody (clone 1A4; Sigma-Aldrich, C6198; 1:500), and DAPI (Sigma-Aldrich, D9542; 1:300). Sections were washed with PBS containing Tween-20 between incubation steps and mounted using Dako Fluorescence Mounting Medium (Agilent). Images were acquired using a Nikon AX confocal microscope at the Augusta University Imaging Core.

Morphometric measurements were performed using ImageJ. Medial wall thickness was calculated as the mean of eight radial measurements across the α-smooth muscle actin-positive media of each vessel. Lumen area was defined as the area enclosed by the inner boundary of the media, and medial wall area was calculated as the area enclosed by the outer medial boundary minus the lumen area. Measurements of the DA and ascending aorta were paired within each tissue section. Accordingly, the ascending aorta served as the comparator for morphometric analyses, whereas transcriptomic comparisons used pooled cells from the ascending and descending aorta.

### CosMx Spatial Molecular Image (SMI) tissue preparation and profiling

CosMx SMI profiling was performed in the separate cohort of mice described above, with one tissue section analyzed per animal. Mouse tissues were fixed, cryoprotected, embedded, and sectioned as described above for immunofluorescence and wall morphometry. The human specimen was fixed in 4% paraformaldehyde overnight, paraffin-embedded, and sectioned at 5 μm. Additional tissue preparation details are provided in the Supplemental Methods.

Spatial transcriptomic profiling of tissue sections was performed using the CosMx Spatial Molecular Imager (Bruker Spatial Biology) with the RNA assay (chemistry version 6). Mouse sections were assayed using the 965-plex Mouse Universal Cell Characterization Panel, with the standard 50-target add-on module replaced by a 15-target custom add-on (Supplemental Table S2). The human section was assayed with the 1,000-plex Human Universal Cell Characterization Panel with its standard add-on module. The mouse and human panels shared 843 gene symbols (842 after excluding hemoglobin), and cross-species comparisons were restricted to these shared genes. The 20 mouse sections were acquired across seven imaging runs, yielding 3,910,156 segmented cells, whereas the human section was acquired in a single run, yielding 102,033 segmented cells. Acquisition and quality-control metrics for each run are provided in Supplemental Table S3.

### Data processing and spatial annotation

Cell-level quality control of mouse sections was performed separately for each imaging run. Depending on the run, a minimum threshold of 20–50 total transcripts per cell was applied, together with criteria for negative-probe counts, transcript complexity or number of detected genes, and cell area, consistent with AtoMx SIP defaults. The criteria applied to each run are provided in Supplemental Table S3. Human cells were retained if they had at least 50 total counts across all 1,207 probes, including target, negative-control, and system-control probes, with no more than 5% of total counts derived from negative-control probes.

The DA, ascending aorta, and descending aorta were manually outlined in each mouse tissue section using a custom browser-based annotation tool, yielding 60 initial vessel regions. Each region was subsequently redefined using a single geometric expansion rule applied identically across vessels and sections, independently of gene expression (Supplemental Methods). Cells added through this expansion were subjected to the same run-specific quality-control criteria as cells within the original vessel outlines. This procedure yielded 48,940 annotated vessel-wall cells with expression data. Subsequently, 96 luminal objects, all from P2.5 sections, were removed using a geometric rule (Supplemental Methods). An additional 22 cells remained unassigned after lineage classification and were excluded, leaving 48,822 cells for downstream analyses.

Cells were grouped by developmental stage and clustered using Seurat v5 with SCTransform normalization.^37,38^ Clusters were annotated based on canonical lineage marker expression. Clusters characterized by hemoglobin transcripts, consistent with erythroid cells or ambient RNA contamination, and one cardiomyocyte cluster were excluded. Unassigned cells, cells added through geometric expansion, and all cells from the two P2.5 imaging runs were classified using a multinomial logistic regression model trained on marker-supported cells from the E14.5 to P0.5 runs with 54 panel-verified markers (Supplemental Table S2) and validated using held-out blocks (Supplemental Methods). Because *Notch3* and *Pdgfrb* expression did not reliably distinguish SMCs from pericytes, these populations were combined into a single mural cell population. Human cells were clustered and annotated using the same general approach, yielding 14 categories. Of 69,390 human cells passing quality control, 40,765 were assigned to the 11 cell populations included in the reported analyses. Two-dimensional embeddings were generated using Harmony integration by imaging run, followed by uniform manifold approximation and projection (UMAP) (Supplemental Methods).^39,40^

### Cell composition, mural programs and gene and pathway analyses

Expression values used for visualization or direct comparison between genes were expressed as log1p-transformed counts per 10,000 (CP10k). Signaling-module scores were calculated separately using counts normalized to a total of 100 per cell, followed by log1p transformation. Vessel contrasts were expressed as AUC − 0.5, where AUC = U/(n_DA × n_Ao), and were calculated within each section by comparing DA cells with pooled aortic cells. A value of zero indicated no directional separation, whereas positive values indicated higher values in the DA. Comparisons of gene expression or detection rates across developmental stages were depth-matched by binomial downsampling of each cell’s counts to a common depth (Supplemental Methods).

Cell population composition was defined as the percentage of annotated cells belonging to each population within each vessel, with section-level percentages used as the unit of analysis. Within each developmental stage, DA and aortic proportions were compared using paired *t* test of within-section differences, with residual variance pooled across the five developmental stages (15 degrees of freedom). Homogeneity of variance across stages was assessed using the Brown– Forsythe test. Comparisons between developmental stages were performed using Welch’s *t* tests on section-level percentages.

Mural cells were scored for five transcriptional programs: contractile, contractile-remodeling, proliferative, matricellular-remodeling, and matrix-remodeling. Each program was defined by a prespecified marker-gene set (Supplemental Table S2). The raw score for each program was calculated as the mean log1p CP10k expression of its marker genes. Scores were then corrected for total transcript count. For each program, a linear relationship constrained through the origin was fitted between the raw program score and log-transformed total transcript counts, centered on the mean of DA and ascending aortic cells, using only these two vessel populations. The fitted depth-dependent component was then subtracted from the scores of cells across all three vessels. Corrected scores were standardized within each section relative to ascending aortic mural cells using a robust *z* score, calculated as the difference between the cell score and the reference median divided by the median absolute deviation of the reference cells. Each mural cell was assigned to the program with the highest standardized score, resulting in one predominant program label per cell. Program composition was expressed as the percentage of mural cells assigned to each program within each vessel and section.

Contractile-associated gene expression was further summarized using an eight-gene composite score (Supplemental Table S2) calculated from within-cell gene ranks. For each mural cell with at least 250 total counts, all 965 panel genes were ranked by transcript count using midranks for ties. The contractile score was calculated as the mean percentile rank of the eight genes on a scale from 0 to 100 (Supplemental Methods). Section-level mean scores were calculated separately for each vessel and compared within sections. Signaling-module comparisons were repeated within each mural cell program, comparing cells with the same program label, to assess whether vessel-specific differences persisted after accounting for differences in program composition (Supplemental Methods).

Sixteen signaling modules comprising 221 panel genes (Supplemental Table S2) were scored in each cell as the mean expression of module genes minus the mean expression of expression-matched control genes (Supplemental Methods). Within each section and cell population, the DA was compared with the pooled aorta using AUC-based contrasts. Macrophage abundance met the prespecified inclusion threshold in only 13 of 20 sections; therefore, macrophage results were displayed descriptively without significance testing.

Stage-specific cell-cycle-associated transcription was evaluated using a wall-wide cell-cycle score, defined as the mean of the within-section DA–aorta differences in cell-cycle module expression across endothelial cells, mural cells, and adventitial fibroblasts. This composite score was defined after inspection of the lineage-specific results and was tested once per developmental stage, with multiple-testing correction across the five stages. Lineage-specific values were reported as components of the composite score and were not tested separately.

For gene-level analyses, DA and pooled aortic cells were compared within each developmental stage across all five cell populations using two-sided Mann-Whitney tests. AUC and log2 fold change, calculated using the *scanpy* rank_genes_groups approach, were reported as effect-size measures. Hemoglobin genes were excluded from all DA–aorta comparisons, leaving 963 of the 965 panel targets. Significance required a Benjamini-Hochberg-adjusted *q* value below 0.05 within each stage across the 963 targets and an absolute log2 fold change of at least 0.5.^41^ Because this screening analysis treated individual cells as observations, it was used only to identify and rank candidate genes. Vessel-specific differences reported as inferential findings in the Results were tested using tissue sections as the unit of analysis. Per-section analyses were restricted to genes detected in at least 5% of cells within the population tested, with Benjamini-Hochberg correction applied across eligible genes.

Fibroblast extracellular matrix genes were identified from the panel-wide, section-level screen and are reported with their panel-wide *q* values. Genes comprising the adventitial insulin-like growth factor axis and endothelial integrins were individually evaluated using section-level AUC contrasts. Each gene set was also compared with a background set of 300 panel genes scored using the same approach to distinguish an overall shift in gene-set expression from differences in the relative composition of the set.

To identify genes for which DA–aorta differences changed across development, a section-level contrast (*e* = AUC − 0.5) was calculated for each detected gene. One-way analysis of variance across developmental stages was then performed using the tissue section as the biological replicate. A gene was described as diverging postnatally when the absolute mean contrast was below 0.1 at every prenatal stage and at least 0.1 at a postnatal stage. Because endothelial cells and adventitial fibroblasts within the vessel wall could contain transcripts originating from adjacent mural cells, expression of each gene was first adjusted for a per-cell mural score, defined as the rank-transformed mean expression of *Myh11* and *Actg2*. Both screens were then performed on the resulting residuals. *Myh11* and *Actg2* were excluded from the reported results because they were used to define the mural score.

### External dataset analyses

To determine whether selected mural, endothelial, and fibroblast differences identified by spatial profiling were also observed in an independent dataset, we reanalyzed published single-cell RNA sequencing data from the mouse DA and descending aorta at E18.5, P0.5, and P5 (GEO accession GSE188202; Supplemental Table S1).^20^ Count data were processed using Seurat v5. Smooth muscle cells, endothelial cells, and fibroblasts were identified using the same canonical markers applied to the spatial dataset. For each gene, expression was averaged within each cell population, vessel, and developmental stage and expressed as log2(CPM + 1). The DA–aorta difference was then calculated. For the cell-cycle entries in Supplemental Table S1, the reported value represents the median DA–aorta difference across nine proliferation-associated genes (*Mki67*, *Top2a*, *Cenpf*, *Cdk1*, *Ube2c*, *Tpx2*, *Birc5*, *Ccna2*, and *Nusap1*). The contractile and matrix-remodeling composite entries were calculated in the same way across the corresponding gene sets in Supplemental Table S2. Because each library in the published dataset comprised pooled samples from littermates, with only one library available per vessel at each developmental stage, independent biological replicates were unavailable for statistical testing. Accordingly, these analyses were considered descriptive and were used only to assess whether DA–aorta differences showed directional consistency with those observed in the spatial data.

### Statistical analysis

For wall morphometry, individual animals served as the biological replicates. Each morphometric measure was analyzed in GraphPad Prism using ordinary one-way analysis of variance (ANOVA) across the 10 developmental stage-by-vessel groups, followed by Tukey’s multiple-comparisons test. Of these comparisons, the five prespecified within-stage comparisons between the DA and ascending aorta are reported with their multiplicity-adjusted *P* values.

For spatial gene profiling, each tissue section represented one animal and served as the unit of analysis for section-level comparisons. Vessel comparisons were paired within each section, and, unless otherwise specified, section-level differences were evaluated using two-sided one-sample *t* test against the null hypothesis of no difference. For gene-set background comparisons, genes rather than sections served as observations, and each gene set was compared with its background using a two-sided Mann-Whitney test. The pooled gene-level screen and one-versus-rest marker analyses used for cell-type annotation treated individual cells as observations and were used for gene ranking and candidate identification rather than statistical inference.

Multiple-testing correction families were defined separately for each analysis. For cell-composition analyses, DA–aorta comparisons within developmental stages and comparisons across developmental stages constituted separate families, with Holm correction applied within each cell population or mural program. Planned comparisons across developmental stages, such as those involving the contractile rank score, were corrected across the five stages using the Benjamini-Hochberg procedure.

Gene-level and module-level screens were corrected using the Benjamini-Hochberg procedure across all genes or modules included in the respective screen. For stage-resolved follow-up analyses of a module or contrast that met the significance criterion in its primary screen, multiplicity correction was applied across the five developmental stages. Results outside these defined testing families were considered descriptive and are presented without significance indicators. Exact *P* and *q* values are reported.

Analyses were performed in R version 4.6.0 using Seurat and ggplot2, and in Python version 3.11.15 using Scanpy, AnnData, scikit-learn, SciPy, statsmodels, NumPy, pandas, and diptest. Individual package versions are provided in the Supplemental Methods. Claude (Anthropic) was used to assist with the development of analysis code. The final analysis code was reviewed and validated by the authors.

## Acknowledgments

We thank the members of the Vascular Biology Center at Augusta University for their support.

## Author Contributions

C.S. and J.S. conceived and designed the study. C.S., I.D., and J.S. performed the experiments. J.D.S. provided human DA tissue. J.R., E.L.S., and B.K.S. contributed to the study design and scientific direction. M.Z. performed spatial transcriptomic profiling. C.S., A. Sinha, and A. Sharma performed the bioinformatic and computational analyses. C.S. and J.S. interpreted the findings, prepared the figures, and wrote the manuscript. All authors reviewed and edited the manuscript and approved the final version.

## SUPPLEMENTAL MATERIAL

### SUPPLEMENTAL METHODS

#### Tissue preparation

Pregnant dams were euthanized by isoflurane inhalation, and embryos were immediately collected, separated from the placenta and extraembryonic membranes, and euthanized by hypothermia in ice-cold Dulbecco’s phosphate-buffered saline (DPBS). Postnatal pups were euthanized by an overdose of inhaled isoflurane. Embryos and pups were fixed in 4% paraformaldehyde for 2 hours at 4°C, washed with DPBS, and cryoprotected overnight at 4°C in 30% sucrose prepared in phosphate-buffered saline (PBS). Tissues were then embedded in OCT compound (Sakura Tissue-Tek), frozen on dry ice, and stored at −80°C. Cryosections (5 μm) were prepared using a Leica CM1950 cryostat with the chamber temperature set to −20°C, mounted on Leica BOND PLUS slides (Leica Biosystems, S21.2113.A), and stored at −80°C until further processing. For CosMx profiling, slides were transferred directly from −80°C into prechilled 10% neutral-buffered formalin and maintained on ice for 30 minutes. Fixation was then continued at 4°C for a total fixation time of 2 hours at the genomics core.

Human tissue was transferred directly from the operating room and fixed overnight in 4% paraformaldehyde. Three tissue segments were embedded in OCT compound, and two were paraffin-embedded. Sections were cut at 5 μm, and one section from the paraffin-embedded tissue was used for CosMx profiling.

#### Imaging and quality control

Mouse and human sections were processed according to the manufacturer’s protocols for fresh-frozen (mouse) and formalin-fixed paraffin-embedded (human) tissue (CosMx SMI Manual Slide Preparation for RNA Assays, MAN-10184). Briefly, after antigen retrieval, slides were rinsed in DEPC-treated water, washed for 3 minutes in 100% ethanol, digested mildly with proteinase K, rinsed, incubated with fiducial solution, and fixed in 10% neutral-buffered formalin. Samples were then quenched and blocked with 100 mM NHS-acetate in CosMx NHS-acetate buffer. The in situ hybridization probe mix (1 nM probes in DEPC-treated water with CosMx Buffer R and RNase inhibitor) was applied under coverslips and hybridized at 37 °C for 16 hours. Slides were washed, stained with DAPI in blocking buffer, incubated for 1 hour with the morphology antibodies, washed three times in 1X PBS, and assembled into CosMx flow cells. On the instrument, fields of view were placed on the tissue, cell morphology was imaged before RNA readout, and eight Z-stack images (0.8 μm step) were acquired per field of view in each reporter cycle; reporter fluorophores were cleaved by UV illumination and removed with Strip Wash Buffer between cycles, and the hybridization and imaging cycle was repeated through all reporter pools and multiple times to increase detection sensitivity. Data were transferred to the AtoMx Spatial Informatics Platform, and post-run quality controls confirmed the hybridization rate and cell segmentation of each field of view. Imaging was performed using two CosMx Spatial Molecular Imager (SMI) instruments running software version 2.2.0.106, with individual fields of view (FOVs) measuring approximately 0.51 × 0.51 mm. The mouse panel contained 965 gene targets, 10 negative-control probes, one additional negative-control probe, and 184 system-control probes. The human panel contained 1,000 gene targets, 10 negative-control probes, and 197 system-control probes. The 20 mouse sections encompassed 2,862 FOVs, whereas the human section encompassed 99 FOVs.

Cell segmentation was performed using the AtoMx Spatial Informatics Platform (version 2.3.2) based on five morphology channels: DAPI, PanCK, CD45, CD68, and CD298/B2M for mouse sections, and DAPI, PanCK, CD45, CK8/18, and a membrane marker for the human section. Gene-expression matrices, cell metadata, segmentation polygons, FOV positions, and transcript coordinates were exported from AtoMx. Run-specific quality-control metrics, including the number of FOVs and segmented cells, median transcript counts and detected genes per cell, mean negative-probe proportion, and median cell area, are provided in Supplemental Table S3.

#### Spatial annotation and cell filtering

Vessel region boundaries were expanded using a single-pass geometric rule. Candidate cells were assigned to a vessel region if their centroids were within 15 μm of the centroid of any cell inside the original manually defined boundary. Cells meeting the criteria for multiple regions were assigned to the nearest region to prevent duplicate assignments. The expansion was noniterative, and newly assigned cells were not used to identify additional candidates. This procedure identified 6,240 candidate cells across the six original imaging runs and 312 in the second P2.5 run, with transcript counts obtained from each run’s complete flat-file export. All candidate cells were subjected to the quality-control criteria of their respective imaging runs (Supplemental Table S3), resulting in the exclusion of 1,186 cells. Thus, geometric expansion added 5,366 cells to the 43,574 cells within the original vessel boundaries, yielding 48,940 annotated vessel-wall cells with expression data. Of these, 75 cells remained unassigned after lineage classification, including 53 among the luminal objects described below.

At P2.5, the open aortic lumens within the manually defined vessel boundaries contained segmented objects with very few transcript counts, consistent with blood cells. Most of these objects were detected in the second imaging run. To identify and remove these objects, vessel lumens were defined geometrically. Cells within each vessel were mapped onto an 8-μm grid, and small gaps in the ring formed by vessel-wall cells were closed using binary morphological closing with a square structuring element with a radius of three grid cells (7 by 7 cells, 56 μm). The enclosed region was then filled, and the area not covered by vessel-wall cells was designated as the lumen. Only cells with transcript counts of at least 25% of the section’s median transcript count were used to construct the vessel-wall ring, preventing low-count objects from contributing to lumen delineation. Cells were excluded only if they were located within the geometrically defined lumen and had transcript counts below this 25% threshold.

This procedure identified 96 of the 11,501 P2.5 cells with expression data, including 77 cells from one section of the second imaging run. These 96 objects were removed from the master data tables used for all subsequent analyses, leaving 48,844 cells. Of these, 22 cells remained unassigned after lineage classification and therefore lacked population labels. Consequently, 48,822 cells were included in cell-composition analyses and all analyses stratified by cell population.

#### Clustering and cell typing

Cell clustering was performed using SCTransform normalization of the NanoString assay (clip.range = c(-10, 10)). Clusters were identified using 30 principal components, with clustering resolutions of 0.5 for mouse sections and 0.8 for the human section. The lineage classifier was trained using 54 marker genes (Supplemental Table S2), with log1p-transformed counts per 10,000 (CP10k) standardized within each developmental stage. Sections were standardized within their respective imaging runs before pooling. Classifier performance was evaluated using leave-one-stage-by-run-block-out validation.

#### Embeddings

The vessel-wall embedding (Supplemental Figure S1B) was generated using 48,822 analyzed cells across all 20 mouse sections. Transcript counts were normalized to a total of 100 per cell and log1p-transformed. Fifty principal components were calculated, and the first 30 were corrected for imaging-run effects using Harmony. A 15-nearest-neighbor graph was constructed from the corrected components and visualized using uniform manifold approximation and projection (UMAP; min_dist = 0.35, fixed random seed). The mural-program embedding (Supplemental Figure S1E) was generated using the same procedure for the 33,710 mural cells assigned to transcriptional programs, with 30 principal components corrected for imaging-run effects.

#### Depth matching

Transcript counts varied across cells, potentially affecting gene-expression levels and detection rates. To account for differences in transcript capture depth when comparing expression levels or detection rates across developmental stages, cells with fewer than 250 total transcripts were excluded, and counts in the remaining cells were randomly downsampled to an expected depth of 250 transcripts per cell using binomial downsampling with a fixed random seed to ensure reproducibility.

#### Program scores

Each mural cell was assigned a score for each transcriptional program. Raw program scores were calculated as the mean log1p-transformed CP10k of the corresponding marker genes (Supplemental Table S2). To account for differences in total transcript counts, a linear relationship constrained through the origin was fitted between each raw program score and log-transformed total transcript counts, centered on the mean of DA and ascending aortic cells. The model was fitted using DA and ascending aortic cells across all developmental stages, and the estimated depth-dependent component was subtracted from the scores of cells in all three vessels.

Within each section, corrected program scores were standardized relative to ascending aortic mural cells using robust *z* scores, calculated as the difference between each cell’s score and the median of the reference cells, divided by 1.4826 times the median absolute deviation of the reference cells. Thus, a standardized score of zero represented the median score of ascending aortic mural cells within the same section. The descending aorta was not used as the reference for standardization but was included in the pooled aortic comparator for subsequent analyses. Each mural cell was assigned to the transcriptional program with the highest standardized score.

#### Module scores

For each cell, signaling-module scores were calculated as the mean expression of module genes minus the mean expression of expression-matched control genes. Control genes were selected from 25 expression bins, using the entire eligible gene pool within each bin, weighted according to the number of module genes assigned to that bin. Module membership is provided in Supplemental Table S2.

For each mural cell with at least 250 total transcripts, all 965 panel genes were ranked by their within-cell transcript counts, with midranks assigned to tied values. Ranks were converted to percentiles, and the contractile composite score was calculated as the mean percentile rank of eight contractile genes, yielding a score ranging from 0 to 100. This approach follows the rank-based scoring principle used by UCell and depends on the relative ranks of contractile genes among all genes measured within the same cell. Two additional versions of the contractile composite score were calculated using the same cells as sensitivity analyses and deposited with the source data. The first measured detection rate, defined as the mean fraction of the eight genes detected per cell after binomial downsampling to 250 counts. The second measured transcript abundance, defined as the summed counts of the eight genes per 10,000 total counts after excluding hemoglobin transcripts. The detection-based composite showed the same direction of vessel differences at every developmental stage.

A vessel within a section was included in a comparison only if it contained a sufficient number of cells from the population under investigation (Figure 1E). Because each mural program was defined by its own marker genes, modules containing those genes could exhibit elevated scores in the corresponding program by construction. Therefore, for within-program comparisons, module scores were recalculated after excluding genes used to define the program being compared, while the original program assignments were retained. The greatest overlap occurred in the cell-cycle module, for which six of the twelve genes (*Mki67*, *Top2a*, *Birc5*, *Pcna*, *Hmgb2*, and *Tyms*) also defined the proliferative program. All other overlaps involved one to three genes (Supplemental Table S2).

#### Background gene comparisons

Two gene sets, endothelial integrins and the adventitial IGF axis, were evaluated both individually and collectively to determine whether their expression was preferentially enriched in the DA or aorta. For each gene, a single effect value was calculated by averaging the within-section DA–aorta differences in area under the receiver operating characteristic curve (AUC − 0.5) across sections. Positive values indicated higher expression in the DA, whereas negative values indicated higher expression in the aorta. The effect values of genes within each set were compared with those of 300 background genes randomly selected without replacement from the panel using a fixed random seed. Background genes were selected from all panel targets for the endothelial integrin analysis and from targets detected in more than 5 % of cells for the adventitial IGF-axis analysis. Background genes were not matched to the detection rates of the genes under investigation in either analysis. Comparisons were performed using two-sided Mann-Whitney tests. Gray shaded bands in Figure 2G and Supplemental Figure S2E represent the 10th to 90th percentile range of the background genes. Neither gene set showed a systematic shift relative to its respective background distribution.

#### Carry-over correction

To account for potential transcript carry-over from adjacent mural cells, a mural-associated score was calculated as the mean log1p-transformed CP10k expression of *Myh11* and *Actg2*, followed by rank transformation within each section. Each gene’s expression was regressed against this score using a quadratic model, and DA–aorta AUC contrasts were calculated from the resulting residuals. For analyses displaying expression levels rather than vessel contrasts, a level-preserving adjustment was applied: Adjusted expression = observed expression − fitted expression + fitted expression at the median mural score.

#### Gene-level screen

Gene-level log2 fold changes were calculated using the scanpy rank_genes_groups formulation with a pseudocount of 1 × 10. Multiple-testing correction was performed using the Benjamini-Hochberg procedure separately within each developmental stage across the 963 genes remaining after exclusion of hemoglobin genes.

#### Software

Analyses were performed in R version 4.6.0 using Seurat 5.5.1, harmony 2.0.5, uwot 0.2.5, and ggplot2 4.0.3. Python version 3.11.15 was used with Scanpy 1.11.5, AnnData 0.12.19, scikit-learn 1.8.0, SciPy 1.17.1, statsmodels 0.14.6, NumPy 2.4.4, pandas 2.3.3, and diptest 0.11.0.

#### Sensitivity analyses

Several sensitivity analyses were performed to evaluate the robustness of cell annotation, transcriptional programs, and vessel-specific differences. The results are reported in the main text and Supplemental Tables.

#### Cell filtering and composition

The effect of the imaging-run-specific minimum transcript-count threshold was evaluated in the imaging run containing the second E18.5 and P0.5 pairs, for which a region-of-interest export was available. A total of 319 cells that passed AtoMx quality-control flags but were excluded by the transcript-count threshold were restored, assigned lineage identities using the classifier described above, and included in repeated cell-composition analyses. Only one statistical borderline classification changed: the mural-cell difference at P0.5 (*P* = 0.046 in the primary analysis versus *P* = 0.067 after cell restoration). No stage-level mean changed by more than 0.4 percentage points. Complete results are deposited with the source data.

#### Aortic comparators and spatial boundaries

The suitability of pooling ascending and descending aortic cells was evaluated using two complementary approaches. First, mural cells from the two aortic regions were compared directly using section-level AUC contrasts. The DA was also compared with each aortic segment separately using the same section-level contrasts. Second, in each corrected embedding, the regional composition of the 15 nearest neighbors of each cell was assessed. Among the 15 nearest neighbors of ascending-aortic cells, an average 47.2% were descending-aortic cells and 37.0% were ascending-aortic cells in the whole-wall embedding (49.4% and 40.8% in the mural embedding). In contrast, DA cells predominantly had neighbors from the DA (62.8% in the whole-wall embedding and 75.3% in the mural embedding). These findings indicate substantial transcriptional overlap between the two aortic regions, supporting their use as a pooled comparator, while suggesting greater separation between aortic and ductal cells. The manually defined vessel boundaries were independently assessed by examining *Myh11* expression within a 15-μm band immediately outside each original boundary.

#### Cell identity and mural programs

Cell-population identities were evaluated using one-versus-rest Mann-Whitney tests across all 965 panel targets. The structure of the mural compartment was assessed using clustering resolutions ranging from 0.2 to 1.0, Hartigan’s dip tests within developmental stages, two-component model comparisons based on the Bayesian information criterion (BIC), silhouette width, and a panel-wide search for independently identified markers after excluding genes used to define the original program labels.

#### Module scoring, cell composition, carry-over, and screen stability

To assess the sensitivity of module scores to the selection of reference genes, all comparisons were repeated using reference genes randomly sampled from the same expression bins. The conclusions remained unchanged.

To distinguish vessel-specific expression differences from those attributable to cell composition, artificial vessel profiles were constructed in which gene expression within each cell population was held constant across vessels, while cell-population proportions were allowed to vary. Differences between these profiles therefore reflected differences in cell composition alone. Gene-level comparisons were then repeated using tissue sections as the unit of analysis.

To evaluate the correction for potential mural transcript carry-over into endothelial cells, three additional mural markers (*Acta2*, *Tagln*, and *Myl9*) that were not used to construct the correction score were examined after adjustment. Their residual signals in endothelial cells were eliminated, consistent with effective removal of mural-associated transcript contamination.

Finally, to assess the stability of the developmental gene-level screen, the analysis was repeated 20 times using a leave-one-section-out approach. Genes were retained only if they remained statistically significant in all 20 iterations.

### SUPPLEMENTAL FIGURE LEGENDS

**Supplemental Figure S1.**
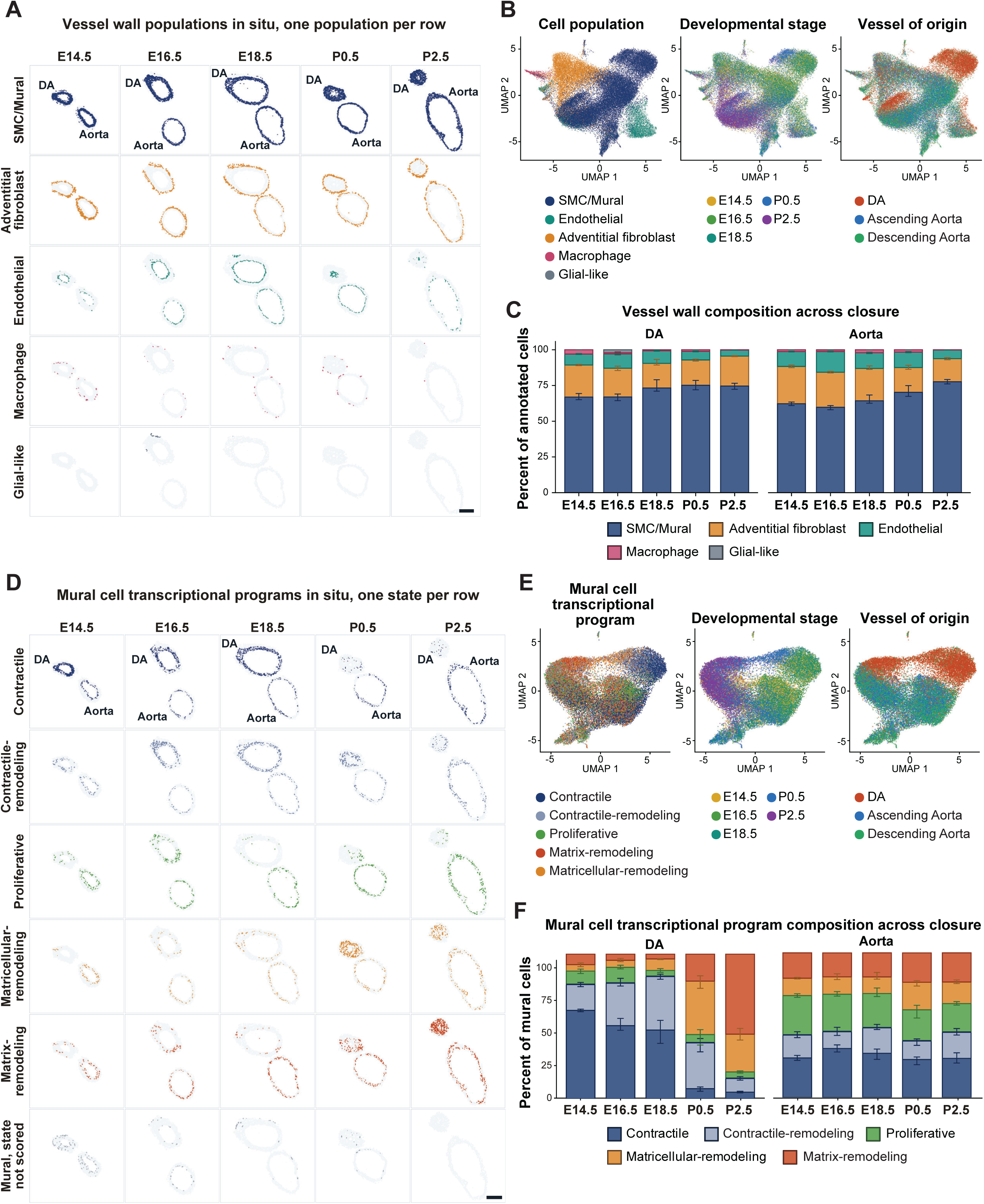
Cell population composition, mural cell transcriptional programs, and spatial distributions during ductus arteriosus development. (A) Spatial distributions of individual cell populations across developmental stages, with the ascending aorta shown as the representative aortic segment. Scale bar, 200 μm. (B) Uniform manifold approximation and projection (UMAP) embeddings of vessel-wall cells colored by cell population, developmental stage, and vessel of origin. A total of 48,822 cells from 20 tissue sections were included (Supplemental Methods). (C) Relative populations of the five annotated cell populations in the ductus arteriosus (DA) and pooled aorta across developmental stages. Values represent the percentages of annotated cells within each vessel and do not indicate absolute cell numbers. Cell proportions were calculated before glial-like cells were excluded from subsequent transcriptional analyses. Bars represent mean section-level percentages ± SEM (n = 4 sections per stage). (D) Spatial distributions of individual mural cell transcriptional programs across developmental stages. Mural cells without a program assignment are shown in the bottom row. Scale bar, 200 μm. (E) UMAP embeddings of mural cells colored by transcriptional program, developmental stage, and vessel of origin. A total of 33,710 mural cells from the same 20 tissue sections were included. (F) Relative proportions of mural cell transcriptional programs in the DA and pooled aorta across developmental stages. Bars represent mean section-level percentages ± SEM (n = 4 sections per stage). DA, ductus arteriosus; SEM, standard error of the mean; UMAP, uniform manifold approximation and projection.

**Supplemental Figure S2.**
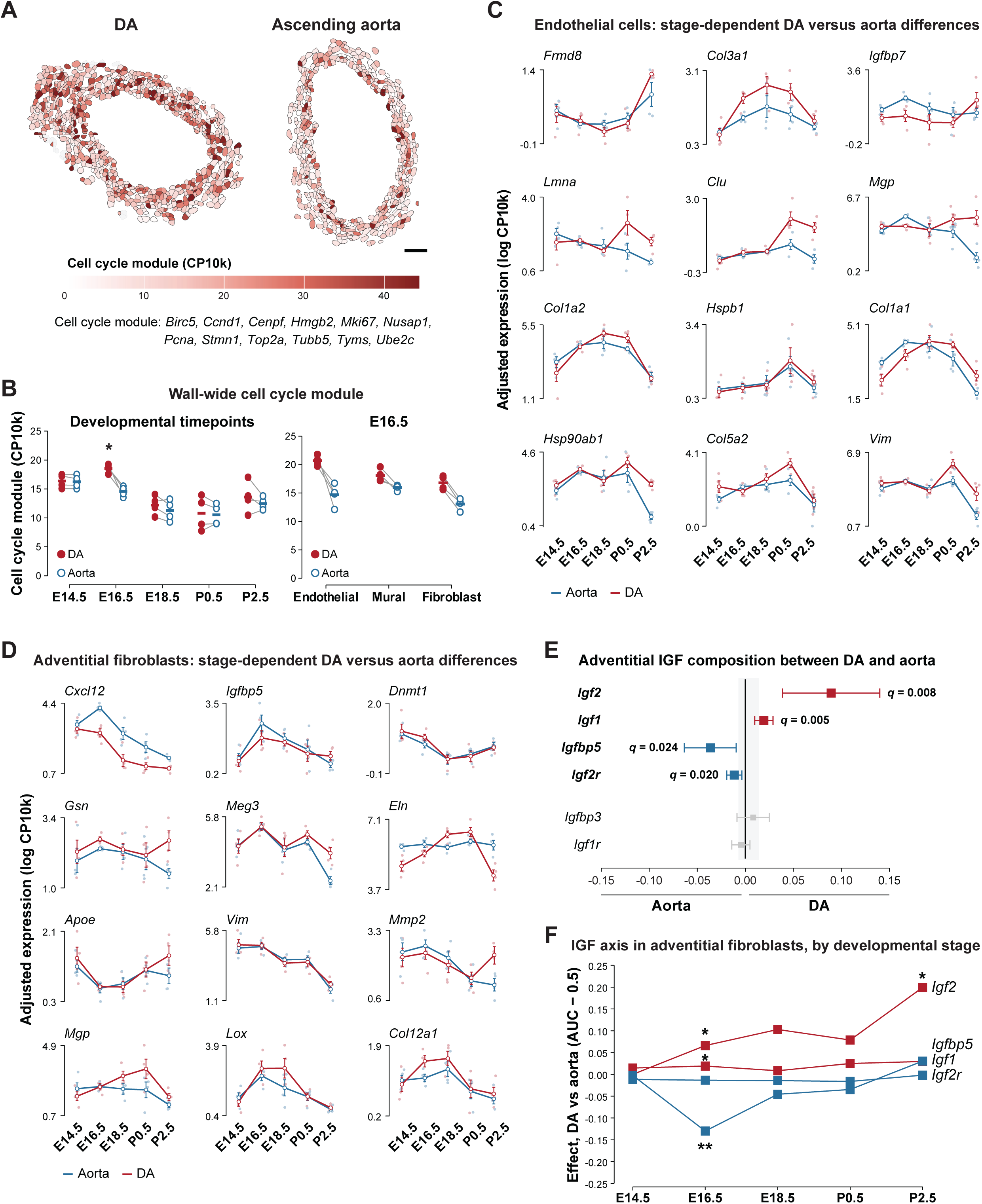
Developmental gene expression in endothelial cells and adventitial fibroblasts of the ductus arteriosus and aorta. **(A)** Spatial distribution of cell-cycle module scores in a representative E16.5 tissue section. Endothelial cells, mural cells, and adventitial fibroblasts are colored according to their cell-cycle module score; macrophages and glial-like cells are shown in gray. Scale bar, 50 μm. **(B)** Wall-wide cell-cycle module scores, calculated as the mean of the endothelial, mural, and adventitial fibroblast scores within each vessel, in the ductus arteriosus (DA) and pooled aorta across developmental stages (left), and cell population– specific scores at E16.5 (right). Points represent individual tissue sections (n = 4 per stage), and bars indicate the mean. \**q* < 0.05, one-sample *t* test of within-section DA-aorta differences with Benjamini-Hochberg correction across the five stages; population-specific values are descriptive. **(C)** Developmental expression profiles of selected endothelial genes in the DA and pooled aorta, expressed as adjusted log counts per 10,000 transcripts (log CP10k). The 12 genes with the smallest vessel-by-stage interaction *P* values are shown, arranged by similarity of their developmental profiles. Red indicates the DA, and blue indicates the aorta. Faint points represent individual tissue sections, whereas open circles and error bars indicate the mean ± SEM (n = 4 sections per stage; 20 sections total). **(D)** Developmental expression profiles of selected adventitial fibroblast genes in the DA and pooled aorta, displayed as described in (C) (n = 20 sections across five developmental stages). **(E)** Vessel contrasts (AUC − 0.5) for insulin-like growth factor (IGF) axis genes between DA and aortic adventitial fibroblasts. Points represent the mean within-section DA–aorta contrast across 20 paired tissue sections, and error bars indicate 95% confidence intervals. Red indicates higher expression in the DA, blue indicates higher expression in the aorta, and gray indicates genes without statistically significant differences. The shaded region represents the 10th to 90th percentile range of the corresponding contrasts for 300 randomly selected panel genes. Adjusted *q* values are shown for genes with statistically significant differences. **(F)** Stage-specific DA-aorta expression contrasts for the IGF axis genes shown in (E). \**q* < 0.05; \*\**q* < 0.01. Ao, aorta; AUC, area under the curve; DA, ductus arteriosus; IGF, insulin-like growth factor; SEM, standard error of the mean.

**Supplemental Figure S3.**
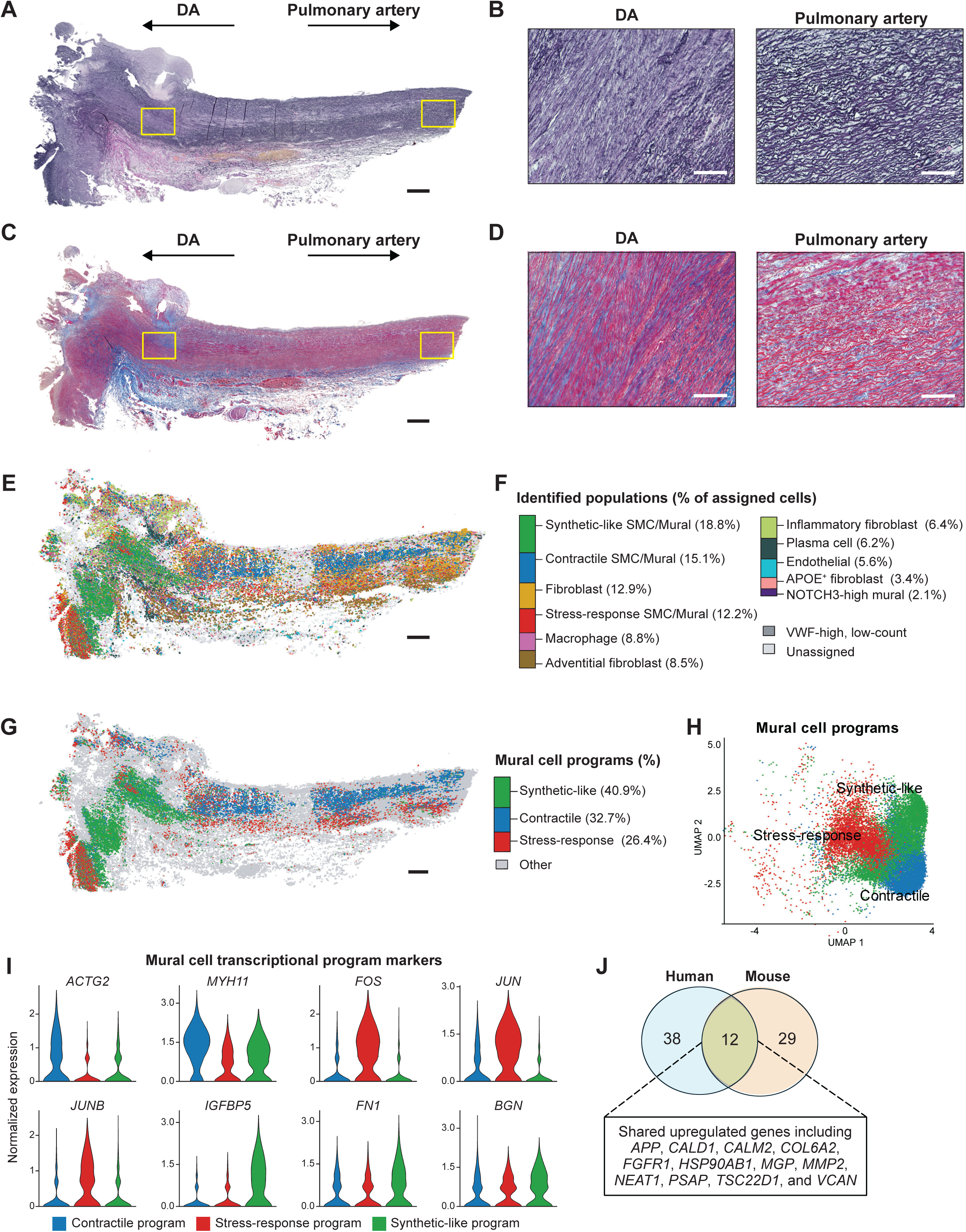
Histology and spatial gene expression. profiles of a human ductus arteriosus and adjacent pulmonary artery. **(A, B)** Elastica van Gieson staining of the human ductus arteriosus (DA) and pulmonary artery. Boxed regions in (A) are shown at higher magnification in (B). Scale bars, 500 μm in (A) and 100 μm in (B). **(C, D)** Masson’s trichrome staining of the same tissue, highlighting collagen in blue. Boxed regions in (C) are shown at higher magnification in (D). Scale bars, 500 μm in (C) and 100 μm in (D). **(E)** Spatial distribution of annotated cell populations. Gray indicates unassigned cells and the VWF-high population (36% of the 69,390 quality-controlled cells in the section). Erythrocytes and hemoglobin-associated low-count cells (5% of quality-controlled cells) are not shown. **(F)** Relative abundance of annotated cell populations, expressed as percentages of assigned cells. **(G)** Spatial distribution of mural cell transcriptional programs. **(H)** Uniform manifold approximation and projection (UMAP) embedding of mural cells colored by transcriptional program. **(I)** Expression of representative marker genes across mural cell transcriptional programs: *MYH11* and *ACTG2* (contractile), *FOS*, *JUN*, and *JUNB* (stress-response), and *IGFBP5*, *FN1*, and *BGN* (synthetic-like). **(J)** Overlap between marker genes of the human synthetic-like mural program and postnatal mouse ductal mural markers, restricted to the 842 genes shared between the human and mouse targeted panels. DA, ductus arteriosus; SMC, smooth muscle cell; UMAP, uniform manifold approximation and projection; VWF, von Willebrand factor.

### SUPPLEMENTAL TABLES

**Supplemental Table S1.** Smooth muscle, endothelial, and fibroblast gene expression patterns in GSE188202 compared with spatial transcript profiling findings. Selected genes and cell-cycle-associated expression programs were examined in smooth muscle cells, endothelial cells, and fibroblasts from the mouse ductus arteriosus (DA) and aorta at E18.5, P0.5, and P5 using the independent single-cell RNA-sequencing dataset GSE188202. Values represent DA–aorta differences in log₂(CPM + 1), ordered as E18.5 → P0.5 → P5. Positive values indicating higher expression in the DA, whereas negative values indicate higher expression in the aorta. Cell-cycle scores represent the median DA–aorta expression difference across *Mki67*, *Top2a*, *Cenpf*, *Cdk1*, *Ube2c*, *Tpx2*, *Birc5*, *Ccna2*, and *Nusap1*. Contractile and matrix-remodeling composites represent the median DA–aorta difference across the contractile composite genes and the matrix-remodeling program genes listed in Supplemental Table S2. Because GSE188202 contains one pooled expression profile per vessel–stage group, these comparisons are descriptive and do not support replicate-level statistical inference.

| **Cell population** | **Gene or program** | **DA–aorta difference (E18.5 → P0.5 → P5)** | **Relationship to spatial finding** |
|---|---|---|---|
| Smooth muscle | *Cald1* | +0.05 → −0.30 → −0.64 | DA enrichment at all five stages; not reproduced after E18.5 |
| Smooth muscle | *Col6a2* | +0.25 → +1.48 → +0.81 | DA enrichment at all five stages |
| Smooth muscle | *Hspb1* | +0.66 → +0.22 → +0.34 | DA enrichment at all five stages |
| Smooth muscle | *Mmp14* | +0.55 → +1.77 → +1.13 | DA enrichment at all five stages |
| Smooth muscle | *Sox9* | +4.62 → +4.75 → +5.63 | DA enrichment at all five stages |
| Smooth muscle | *Tfap2b* | +5.56 → +6.08 → +6.46 | DA enrichment at all five stages |
| Smooth muscle | *Thbs1* | +2.12 → +2.60 → +4.40 | DA enrichment at all five stages |
| Smooth muscle | *Tsc22d1* | +0.99 → +2.26 → +1.63 | DA enrichment at all five stages |
| Smooth muscle | *Tshz2* | +4.00 → +7.13 → +5.90 | DA enrichment at all five stages |
| Smooth muscle | *Cxcl12* | −1.27 → −1.38 → −0.60 | Aorta enrichment at all five stages |
| Smooth muscle | *Itga8* | −1.01 → −1.93 → −1.11 | Aorta enrichment in mural cells at all five stages |
| Smooth muscle | Contractile composite | +0.28 → −0.44 → −0.80 | Postnatal loss of DA contractile enrichment |
| Smooth muscle | *Klf4* | +0.43 → +0.76 → +2.32 | Postnatal DA enrichment |
| Smooth muscle | Matrix-remodeling composite | +0.08 → +1.74 → +1.13 | Postnatal DA enrichment |
| Endothelial | *Vim* | 0.00 → +0.98 → +0.28 | Postnatal DA enrichment |
| Endothelial | Cell cycle | +0.09 → +0.66 → −0.44 | Transient perinatal DA enrichment |
| Fibroblast | *Dcn* | +0.12 → +0.64 → +0.97 | DA-enriched proteoglycan-associated expression |
| Fibroblast | *Lum* | +0.77 → +0.88 → +1.42 | DA-enriched proteoglycan-associated expression |
| Fibroblast | *Mmp2* | +0.49 → +0.68 → +1.04 | Increasing postnatal DA enrichment |
| Fibroblast | *Col1a2* | −0.37 → −0.19 → −0.34 | Persistent aorta-enriched expression |
| Fibroblast | *Col16a1* | −0.24 → −0.35 → −0.57 | Persistent aorta-enriched expression |
| Fibroblast | *Eln* | +0.13 → −0.25 → −0.60 | Postnatal loss of relative DA elastin expression |
| Fibroblast | Cell cycle | +0.55 → −1.23 → −1.36 | Prenatal DA enrichment followed by reversal |

**Supplemental Table S2.** Gene sets used for spatial transcript profiling. This table lists 54 lineage classification markers grouped by cell lineage; marker genes defining five mural cell transcriptional programs; eight genes comprising the contractile composite score; 221 genes assigned across 16 signaling modules; and 15 custom add-on targets that replaced the standard add-on module of the Mouse Universal Cell Characterization Panel. Some genes are shared across multiple mural transcriptional programs.

| **Gene set** | **Lineage or program** | **Gene** |
|---|---|---|
| Lineage classifier marker | Endothelial | *Pecam1* |
| Lineage classifier marker | Endothelial | *Cdh5* |
| Lineage classifier marker | Endothelial | *Kdr* |
| Lineage classifier marker | Endothelial | *Tek* |
| Lineage classifier marker | Endothelial | *Eng* |
| Lineage classifier marker | Endothelial | *Cd34* |
| Lineage classifier marker | Endothelial | *Vwf* |
| Lineage classifier marker | Endothelial | *Esam* |
| Lineage classifier marker | Endothelial | *Ramp2* |
| Lineage classifier marker | Endothelial | *Icam2* |
| Lineage classifier marker | SMC/Mural | *Acta2* |
| Lineage classifier marker | SMC/Mural | *Myh11* |
| Lineage classifier marker | SMC/Mural | *Tagln* |
| Lineage classifier marker | SMC/Mural | *Myocd* |
| Lineage classifier marker | SMC/Mural | *Notch3* |
| Lineage classifier marker | SMC/Mural | *Pdgfrb* |
| Lineage classifier marker | SMC/Mural | *Rgs5* |
| Lineage classifier marker | SMC/Mural | *Actg2* |
| Lineage classifier marker | SMC/Mural | *Tpm2* |
| Lineage classifier marker | Adventitial fibroblast | *Pdgfra* |
| Lineage classifier marker | Adventitial fibroblast | *Col1a1* |
| Lineage classifier marker | Adventitial fibroblast | *Col1a2* |
| Lineage classifier marker | Adventitial fibroblast | *Dcn* |
| Lineage classifier marker | Adventitial fibroblast | *Lum* |
| Lineage classifier marker | Adventitial fibroblast | *Gsn* |
| Lineage classifier marker | Adventitial fibroblast | *Col3a1* |
| Lineage classifier marker | Adventitial fibroblast | *Fbln1* |
| Lineage classifier marker | Adventitial fibroblast | *Mgp* |
| Lineage classifier marker | Adventitial fibroblast | *Col5a1* |
| Lineage classifier marker | Macrophage | *Cd68* |
| Lineage classifier marker | Macrophage | *Csf1r* |
| Lineage classifier marker | Macrophage | *Mrc1* |
| Lineage classifier marker | Macrophage | *Itgam* |
| Lineage classifier marker | Macrophage | *Cd14* |
| Lineage classifier marker | Macrophage | *C1qa* |
| Lineage classifier marker | Macrophage | *C1qb* |
| Lineage classifier marker | Macrophage | *Aif1* |
| Lineage classifier marker | Macrophage | *Tyrobp* |
| Lineage classifier marker | Lymphoid/NK | *Cd3e* |
| Lineage classifier marker | Lymphoid/NK | *Cd3d* |
| Lineage classifier marker | Lymphoid/NK | *Cd2* |
| Lineage classifier marker | Lymphoid/NK | *Nkg7* |
| Lineage classifier marker | Lymphoid/NK | *Gzma* |
| Lineage classifier marker | Lymphoid/NK | *Klrb1c* |
| Lineage classifier marker | Lymphoid/NK | *Cd19* |
| Lineage classifier marker | Lymphoid/NK | *Ms4a1* |
| Lineage classifier marker | Lymphoid/NK | *Il7r* |
| Lineage classifier marker | Lymphoid/NK | *Ccl5* |
| Lineage classifier marker | Lymphoid/NK | *Ptprc* |
| Lineage classifier marker | Other | *S100b* |
| Lineage classifier marker | Other | *Epcam* |
| Lineage classifier marker | Other | *Krt8* |
| Lineage classifier marker | Other | *Krt18* |
| Lineage classifier marker | Other | *Myh6* |
| Mural program | Contractile | *Myh11* |
| Mural program | Contractile | *Tpm2* |
| Mural program | Contractile | *Rgs5* |
| Mural program | Contractile | *Actg2* |
| Mural program | Contractile | *Tagln* |
| Mural program | Contractile | *Acta2* |
| Mural program | Contractile | *Myl9* |
| Mural program | Contractile | *Tpm1* |
| Mural program | Contractile | *Cald1* |
| Mural program | Proliferative | *Tyms* |
| Mural program | Proliferative | *Top2a* |
| Mural program | Proliferative | *Mki67* |
| Mural program | Proliferative | *Pcna* |
| Mural program | Proliferative | *Birc5* |
| Mural program | Proliferative | *Hmgb2* |
| Mural program | Contractile-remodeling | *Tyms* |
| Mural program | Contractile-remodeling | *Tsc22d1* |
| Mural program | Contractile-remodeling | *Mfap5* |
| Mural program | Contractile-remodeling | *Mmp14* |
| Mural program | Contractile-remodeling | *Igf2* |
| Mural program | Contractile-remodeling | *Ptger4* |
| Mural program | Contractile-remodeling | *Thbs1* |
| Mural program | Matricellular-remodeling | *Thbs1* |
| Mural program | Matricellular-remodeling | *Fn1* |
| Mural program | Matricellular-remodeling | *Mgp* |
| Mural program | Matricellular-remodeling | *Tsc22d1* |
| Mural program | Matricellular-remodeling | *Mmp14* |
| Mural program | Matricellular-remodeling | *Bgn* |
| Mural program | Matricellular-remodeling | *Igf2* |
| Mural program | Matrix-remodeling | *Mgp* |
| Mural program | Matrix-remodeling | *Mmp2* |
| Mural program | Matrix-remodeling | *Mmp9* |
| Mural program | Matrix-remodeling | *Mmp14* |
| Mural program | Matrix-remodeling | *Sparcl1* |
| Mural program | Matrix-remodeling | *Mxra8* |
| Mural program | Matrix-remodeling | *Tsc22d1* |
| Mural program | Matrix-remodeling | *Sox4* |
| Mural program | Matrix-remodeling | *Npr1* |
| Contractile composite | Contractile composite | *Acta2* |
| Contractile composite | Contractile composite | *Myh11* |
| Contractile composite | Contractile composite | *Tagln* |
| Contractile composite | Contractile composite | *Myl9* |
| Contractile composite | Contractile composite | *Actg2* |
| Contractile composite | Contractile composite | *Tpm1* |
| Contractile composite | Contractile composite | *Tpm2* |
| Contractile composite | Contractile composite | *Cald1* |
| Signaling module | Cell cycle | *Birc5* |
| Signaling module | Cell cycle | *Ccnd1* |
| Signaling module | Cell cycle | *Cenpf* |
| Signaling module | Cell cycle | *Hmgb2* |
| Signaling module | Cell cycle | *Mki67* |
| Signaling module | Cell cycle | *Nusap1* |
| Signaling module | Cell cycle | *Pcna* |
| Signaling module | Cell cycle | *Stmn1* |
| Signaling module | Cell cycle | *Top2a* |
| Signaling module | Cell cycle | *Tubb5* |
| Signaling module | Cell cycle | *Tyms* |
| Signaling module | Cell cycle | *Ube2c* |
| Signaling module | Chemokine | *Ackr1* |
| Signaling module | Chemokine | *Ackr3* |
| Signaling module | Chemokine | *Ackr4* |
| Signaling module | Chemokine | *Ccl11* |
| Signaling module | Chemokine | *Ccl19* |
| Signaling module | Chemokine | *Ccl2* |
| Signaling module | Chemokine | *Ccl3* |
| Signaling module | Chemokine | *Ccl4* |
| Signaling module | Chemokine | *Ccl5* |
| Signaling module | Chemokine | *Ccl8* |
| Signaling module | Chemokine | *Ccr1* |
| Signaling module | Chemokine | *Ccr2* |
| Signaling module | Chemokine | *Ccr5* |
| Signaling module | Chemokine | *Ccr7* |
| Signaling module | Chemokine | *Cxcl1* |
| Signaling module | Chemokine | *Cxcl10* |
| Signaling module | Chemokine | *Cxcl12* |
| Signaling module | Chemokine | *Cxcl14* |
| Signaling module | Chemokine | *Cxcl16* |
| Signaling module | Chemokine | *Cxcl2* |
| Signaling module | Chemokine | *Cxcl9* |
| Signaling module | Chemokine | *Cxcr3* |
| Signaling module | Chemokine | *Cxcr4* |
| Signaling module | Collagen / ECM core | *Bgn* |
| Signaling module | Collagen / ECM core | *Col11a1* |
| Signaling module | Collagen / ECM core | *Col12a1* |
| Signaling module | Collagen / ECM core | *Col14a1* |
| Signaling module | Collagen / ECM core | *Col15a1* |
| Signaling module | Collagen / ECM core | *Col16a1* |
| Signaling module | Collagen / ECM core | *Col18a1* |
| Signaling module | Collagen / ECM core | *Col1a1* |
| Signaling module | Collagen / ECM core | *Col1a2* |
| Signaling module | Collagen / ECM core | *Col3a1* |
| Signaling module | Collagen / ECM core | *Col4a1* |
| Signaling module | Collagen / ECM core | *Col4a2* |
| Signaling module | Collagen / ECM core | *Col5a1* |
| Signaling module | Collagen / ECM core | *Col5a2* |
| Signaling module | Collagen / ECM core | *Col5a3* |
| Signaling module | Collagen / ECM core | *Col6a1* |
| Signaling module | Collagen / ECM core | *Col6a2* |
| Signaling module | Collagen / ECM core | *Col6a3* |
| Signaling module | Collagen / ECM core | *Col8a1* |
| Signaling module | Collagen / ECM core | *Dcn* |
| Signaling module | Collagen / ECM core | *Eln* |
| Signaling module | Collagen / ECM core | *Fbln1* |
| Signaling module | Collagen / ECM core | *Fn1* |
| Signaling module | Collagen / ECM core | *Lum* |
| Signaling module | Collagen / ECM core | *Mfap5* |
| Signaling module | Collagen / ECM core | *Vcan* |
| Signaling module | Ephrin | *Efna1* |
| Signaling module | Ephrin | *Efna5* |
| Signaling module | Ephrin | *Efnb1* |
| Signaling module | Ephrin | *Efnb2* |
| Signaling module | Ephrin | *Epha2* |
| Signaling module | Ephrin | *Epha3* |
| Signaling module | Ephrin | *Epha4* |
| Signaling module | Ephrin | *Epha7* |
| Signaling module | Ephrin | *Ephb2* |
| Signaling module | Ephrin | *Ephb3* |
| Signaling module | Ephrin | *Ephb4* |
| Signaling module | Ephrin | *Ephb6* |
| Signaling module | FGF | *Fgf1* |
| Signaling module | FGF | *Fgf12* |
| Signaling module | FGF | *Fgf13* |
| Signaling module | FGF | *Fgf18* |
| Signaling module | FGF | *Fgf2* |
| Signaling module | FGF | *Fgf7* |
| Signaling module | FGF | *Fgf9* |
| Signaling module | FGF | *Fgfr1* |
| Signaling module | FGF | *Fgfr2* |
| Signaling module | FGF | *Fgfr3* |
| Signaling module | FGF | *Spry2* |
| Signaling module | FGF | *Spry4* |
| Signaling module | Integrin adhesion | *Itga1* |
| Signaling module | Integrin adhesion | *Itga2* |
| Signaling module | Integrin adhesion | *Itga3* |
| Signaling module | Integrin adhesion | *Itga5* |
| Signaling module | Integrin adhesion | *Itga6* |
| Signaling module | Integrin adhesion | *Itga8* |
| Signaling module | Integrin adhesion | *Itga9* |
| Signaling module | Integrin adhesion | *Itgav* |
| Signaling module | Integrin adhesion | *Itgb1* |
| Signaling module | Integrin adhesion | *Itgb2* |
| Signaling module | Integrin adhesion | *Itgb3* |
| Signaling module | Integrin adhesion | *Itgb5* |
| Signaling module | Integrin adhesion | *Itgb6* |
| Signaling module | Integrin adhesion | *Itgb8* |
| Signaling module | Integrin adhesion | *Ptk2* |
| Signaling module | Interleukin | *Il10* |
| Signaling module | Interleukin | *Il10ra* |
| Signaling module | Interleukin | *Il11* |
| Signaling module | Interleukin | *Il13ra1* |
| Signaling module | Interleukin | *Il15* |
| Signaling module | Interleukin | *Il15ra* |
| Signaling module | Interleukin | *Il17d* |
| Signaling module | Interleukin | *Il17ra* |
| Signaling module | Interleukin | *Il18* |
| Signaling module | Interleukin | *Il1a* |
| Signaling module | Interleukin | *Il1b* |
| Signaling module | Interleukin | *Il1r1* |
| Signaling module | Interleukin | *Il1rn* |
| Signaling module | Interleukin | *Il2rg* |
| Signaling module | Interleukin | *Il33* |
| Signaling module | Interleukin | *Il34* |
| Signaling module | Interleukin | *Il4ra* |
| Signaling module | Interleukin | *Il6* |
| Signaling module | Interleukin | *Il6ra* |
| Signaling module | Interleukin | *Il6st* |
| Signaling module | Interleukin | *Il7r* |
| Signaling module | JAK / STAT | *Jak1* |
| Signaling module | JAK / STAT | *Jak2* |
| Signaling module | JAK / STAT | *Stat1* |
| Signaling module | JAK / STAT | *Stat3* |
| Signaling module | JAK / STAT | *Stat4* |
| Signaling module | JAK / STAT | *Stat5a* |
| Signaling module | JAK / STAT | *Stat5b* |
| Signaling module | JAK / STAT | *Stat6* |
| Signaling module | JAK / STAT | *Tyk2* |
| Signaling module | Matrix proteolysis | *Mmp12* |
| Signaling module | Matrix proteolysis | *Mmp14* |
| Signaling module | Matrix proteolysis | *Mmp19* |
| Signaling module | Matrix proteolysis | *Mmp2* |
| Signaling module | Matrix proteolysis | *Mmp7* |
| Signaling module | Matrix proteolysis | *Mmp9* |
| Signaling module | Matrix proteolysis | *Timp1* |
| Signaling module | Notch | *Dll1* |
| Signaling module | Notch | *Dll4* |
| Signaling module | Notch | *Hey1* |
| Signaling module | Notch | *Jag1* |
| Signaling module | Notch | *Notch1* |
| Signaling module | Notch | *Notch2* |
| Signaling module | Notch | *Notch3* |
| Signaling module | Notch | *Rbpj* |
| Signaling module | PDGF | *Pdgfa* |
| Signaling module | PDGF | *Pdgfb* |
| Signaling module | PDGF | *Pdgfc* |
| Signaling module | PDGF | *Pdgfd* |
| Signaling module | PDGF | *Pdgfra* |
| Signaling module | PDGF | *Pdgfrb* |
| Signaling module | Prostaglandin | *Cyp1b1* |
| Signaling module | Prostaglandin | *Ptgds* |
| Signaling module | Prostaglandin | *Ptger4* |
| Signaling module | Prostaglandin | *Ptges* |
| Signaling module | Prostaglandin | *Ptges2* |
| Signaling module | Prostaglandin | *Ptges3* |
| Signaling module | Prostaglandin | *Ptgis* |
| Signaling module | Prostaglandin | *Ptgs1* |
| Signaling module | Prostaglandin | *Ptgs2* |
| Signaling module | TGF-beta / BMP | *Acvr1* |
| Signaling module | TGF-beta / BMP | *Acvr2a* |
| Signaling module | TGF-beta / BMP | *Acvrl1* |
| Signaling module | TGF-beta / BMP | *Bmp1* |
| Signaling module | TGF-beta / BMP | *Bmp2* |
| Signaling module | TGF-beta / BMP | *Bmp4* |
| Signaling module | TGF-beta / BMP | *Bmp5* |
| Signaling module | TGF-beta / BMP | *Bmp7* |
| Signaling module | TGF-beta / BMP | *Bmpr1a* |
| Signaling module | TGF-beta / BMP | *Bmpr2* |
| Signaling module | TGF-beta / BMP | *Eng* |
| Signaling module | TGF-beta / BMP | *Smad2* |
| Signaling module | TGF-beta / BMP | *Smad3* |
| Signaling module | TGF-beta / BMP | *Smad4* |
| Signaling module | TGF-beta / BMP | *Tgfb1* |
| Signaling module | TGF-beta / BMP | *Tgfb2* |
| Signaling module | TGF-beta / BMP | *Tgfb3* |
| Signaling module | TGF-beta / BMP | *Tgfbr1* |
| Signaling module | TGF-beta / BMP | *Tgfbr2* |
| Signaling module | TNF / TLR / NF-kB | *Cd14* |
| Signaling module | TNF / TLR / NF-kB | *Nfkb1* |
| Signaling module | TNF / TLR / NF-kB | *Nfkbia* |
| Signaling module | TNF / TLR / NF-kB | *Rela* |
| Signaling module | TNF / TLR / NF-kB | *Tlr1* |
| Signaling module | TNF / TLR / NF-kB | *Tlr2* |
| Signaling module | TNF / TLR / NF-kB | *Tlr3* |
| Signaling module | TNF / TLR / NF-kB | *Tlr4* |
| Signaling module | TNF / TLR / NF-kB | *Tlr7* |
| Signaling module | TNF / TLR / NF-kB | *Tnf* |
| Signaling module | TNF / TLR / NF-kB | *Tnfaip6* |
| Signaling module | TNF / TLR / NF-kB | *Tnfrsf11b* |
| Signaling module | TNF / TLR / NF-kB | *Tnfrsf1a* |
| Signaling module | TNF / TLR / NF-kB | *Tnfrsf1b* |
| Signaling module | VEGF / angiopoietin | *Angpt1* |
| Signaling module | VEGF / angiopoietin | *Angpt2* |
| Signaling module | VEGF / angiopoietin | *Cd34* |
| Signaling module | VEGF / angiopoietin | *Flt1* |
| Signaling module | VEGF / angiopoietin | *Kdr* |
| Signaling module | VEGF / angiopoietin | *Tek* |
| Signaling module | VEGF / angiopoietin | *Tie1* |
| Signaling module | VEGF / angiopoietin | *Vegfa* |
| Signaling module | VEGF / angiopoietin | *Vegfb* |
| Signaling module | VEGF / angiopoietin | *Vegfc* |
| Signaling module | VEGF / angiopoietin | *Vegfd* |
| Signaling module | Wnt | *Apc* |
| Signaling module | Wnt | *Ctnnb1* |
| Signaling module | Wnt | *Fzd1* |
| Signaling module | Wnt | *Fzd3* |
| Signaling module | Wnt | *Fzd4* |
| Signaling module | Wnt | *Fzd5* |
| Signaling module | Wnt | *Fzd6* |
| Signaling module | Wnt | *Fzd7* |
| Signaling module | Wnt | *Fzd8* |
| Signaling module | Wnt | *Rspo3* |
| Signaling module | Wnt | *Tcf7* |
| Signaling module | Wnt | *Wif1* |
| Signaling module | Wnt | *Wnt11* |
| Signaling module | Wnt | *Wnt5a* |
| Signaling module | Wnt | *Wnt5b* |
| Signaling module | Wnt | *Wnt7b* |
| Signaling module | Wnt | *Wnt9a* |
| Custom add-on target | Custom add-on | *Bmp10* |
| Custom add-on target | Custom add-on | *Egr1* |
| Custom add-on target | Custom add-on | *Eln* |
| Custom add-on target | Custom add-on | *Fbln1* |
| Custom add-on target | Custom add-on | *Gdf2* |
| Custom add-on target | Custom add-on | *Has2* |
| Custom add-on target | Custom add-on | *Itgb3* |
| Custom add-on target | Custom add-on | *Lox* |
| Custom add-on target | Custom add-on | *Myocd* |
| Custom add-on target | Custom add-on | *Prdm6* |
| Custom add-on target | Custom add-on | *S1pr1* |
| Custom add-on target | Custom add-on | *S1pr2* |
| Custom add-on target | Custom add-on | *S1pr3* |
| Custom add-on target | Custom add-on | *Sphk1* |
| Custom add-on target | Custom add-on | *Tfap2b* |

**Supplemental Table S3.** Imaging acquisition and quality control metrics for CosMx spatial transcript profiling. Each row represents an individual CosMx imaging run (seven mouse runs and one human run). The numbers of fields of view and segmented cells, together with per-cell summary metrics, were obtained from the AtoMx export. Cell-level quality control (QC) criteria are reported separately for each run. For mouse samples, the final three columns indicate the number of spatially annotated vessel-wall cells, the number of luminal objects excluded at P2.5, and the number of cells remaining after geometric and luminal-object filtering, before exclusion of 22 lineage-unassigned cells. For mouse samples, the final three columns indicate the number of spatially annotated vessel-wall cells, the number of luminal objects excluded at P2.5, and the number of cells remaining after geometric and luminal-object filtering, before exclusion of 22 lineage-unassigned cells. The human sample was analyzed as a whole section, for which vessel-wall annotation and luminal-object exclusion were not applicable (NA). Following QC, 69,390 human cells were retained, of which 40,765 were included in the 11 reported cell populations. An additional 3,767 erythrocytes and hemoglobin-associated low-count cells were excluded from the figures.

| **Species and developmental stage** | **Fields of view** | **Segmented cells** | **Median counts per cell** | **Median genes per cell** | **Mean negative-probe proportion (%)** | **Median cell area (µm<sup>2</sup>)** | **Cell-level QC criteria** | **Annotated vessel-wall cells** | **Luminal objects excluded** | Cells retained after filtering |
|---|---|---|---|---|---|---|---|---|---|---|
| mouse, E14.5 | 323 | 536,475 | 180 | 99 | 0.587 | 87.5 | min 20 total counts, with the negative-probe proportion, complexity and cell-area criteria at the instrument's default settings | 7,260 | 0 | 7,260 |
| mouse, E16.5 | 374 | 753,205 | 472 | 154 | 0.112 | 72.4 | min 50 counts, min 50 genes, negative-probe fraction 10% or less, area at or below run 99th percentile | 9,469 | 0 | 9,469 |
| mouse, E18.5 | 363 | 343,521 | 560 | 224 | 0.075 | 96.5 | min 20 total counts, with the negative-probe proportion, complexity and cell-area criteria at the instrument's default settings | 3,776 | 0 | 3,776 |
| mouse, E18.5, P0.5 | 623 | 884,631 | 326 | 130 | 0.148 | 81.8 | min 50 counts, min 50 genes, negative-probe fraction 10% or less, area at or below run 99th percentile | 11,680 | 0 | 11,680 |
| mouse, P0.5 | 302 | 404,049 | 384 | 173 | 0.112 | 76.6 | min 20 total counts, with the negative-probe proportion, complexity and cell-area criteria at the instrument's default settings | 5,254 | 0 | 5,254 |
| mouse, P2.5 (run 1) | 571 | 550,170 | 192 | 115 | 0.527 | 85.2 | min 50 counts, min 50 genes, negative-probe fraction 10% or less, area at or below run 99th percentile | 5,765 | 3 | 5,762 |
| mouse, P2.5 (run 2) | 306 | 438,105 | 209 | 135 | 0.287 | 62.6 | min 20 total counts, with the negative-probe proportion, complexity and cell-area criteria at the instrument's default settings | 5,736 | 93 | 5,643 |
| human, 6 months | 99 | 102,033 | 92 | 53 | 0.206 | 74.4 | min 50 counts over all 1,207 probes, 5% or less negative-probe counts | NA | NA | 69,390 |

NA, not applicable; QC, quality control.

